# Refining the Serine Protease Autotransporters of Enterobacteriaceae (SPATE) gene detection in Enteroaggregative Escherichia coli genomes uncovers differential SPATE distribution by phylogeny

**DOI:** 10.64898/2026.04.16.715897

**Authors:** Rotimi A. Dada, Ayorinde O. Afolayan, Olufunmilayo A. Adewuyi, Babajide A. Tytler, Busayo O. Olayinka, Nicholas R. Thomson, Iruka N. Okeke

## Abstract

**Background:** Enteroaggregative *Escherichia coli* (EAEC) are a heterogenous pathotype, implicated in acute and persistent diarrhoea especially in developing countries. Serine Protease Autotransporters of Enterobacteriaceae (SPATEs) are Type V Secretory System trypsin-like proteases repeatedly reported from EAEC. This study aimed to determine SPATE encoding-gene prevalence among EAEC and their association with diarrhoea. We screened 881 EAEC genomes from four recent epidemiological studies in Nigeria for 23 SPATE-encoding genes, initially using ARIBA and the Virulencefinder database.

**Results:** Initial screening inflated SPATE gene content, particularly in genomes with multiple SPATEs, due to cross detection of highly similar sequences and other artefacts. We developed and validated refined methodology, which detected 478 of 1,156 original SPATE calls and also identified SPATE miscalls from previous datasets in the literature. The most prevalent SPATE-encoding gene in our EAEC collection was *sepA* 297(33.71%), closely followed by *sat* 360 (29.74%). *pic,* encoding a SPATE with mucinase activity, was found in 65 (7.4%) genomes and associated with diarrhoea (*p*=0.00004). EAEC strains belonging to *E. coli* phylogroups A, B1 or C carried, on average, one SPATE gene per genome while >1 was typically detected in phylogroup B2 EAEC. Other EAEC carried few or no SPATE genes.

**Conclusions:** Our study shows that multifunctional genome analysis tools may have to be refined for certain gene families to avoid overestimation. SPATEs are not as prevalent as previously thought but they remain common among EAEC, particularly among phylogroup A, B1, B2 and C, pointing to the possibility that they make lineage-specific contributions to disease.

## Background

Enteroaggregative *Escherichia coli* (EAEC) are a major cause of childhood, visiting travellers’ and other diarrhoea, especially in developing countries [1,2,3]. In developing countries, EAEC has also been linked with malnutrition and growth retardation in children who are at greater risk of diarrhoea-related morbidity and mortality [4,5]. Serine Protease Autotransporters of Enterobacteriaceae (SPATEs) are type V secretion system (T5SS) or autotransporter proteins important for EAEC virulence. These proteins have multiple functions in EAEC pathogenesis and may function as immunomodulators, proteases, adhesins and toxins [6,7,8,9,10]. SPATEs have been shown to enhance virulence in a rabbit model [11] and have also been proposed as markers that could be used to define EAEC and/or delineate potentially hypervirulent strains [12,13,14].

Like other autotransporters, including IgA1 proteases, subtilin-like serine proteases and Self-Associating Autotransporters (SAAT), all SPATEs have 3 specific domains—a *sec*-dependent signal peptide that targets the protein to and through the inner membrane and translocates the rest of the protein from the cytoplasm to the periplasmic space via the *sec* pathway, a N-terminal passenger domain (formerly called the α-domain) which is the functional secreted part and includes a highly conserved serine protease motif (GDSGS); and lastly a C-terminal (formerly called the β-barrel domain), which acts as a pore-forming domain on the outer membrane and translocate the passenger domain to the outer membrane. Additionally, all SPATEs have a linker region which connects the C-terminal pore-forming domain to the N-terminal passenger domain. In some SPATEs the C-terminal is responsible for cleavage of the passenger domain at the linker, within the pores of the outer membrane allowing for its release from the cell. There are 23 SPATE genes described in the literature, including *sat*, *pet*, *vat*, *sigA*, *hbp*, *tsh*, *pic*, *eatA*, *eaaA*, *eaaC*, *sepA*, *espI*, *epeA*, *espP*, espC, *tleA*, *tagB*, *tagC sha*, *boa*, *picU*, *rpeA* and *crc1*. *eaaA* and *eaaC* have been reported to be usually associated with non-pathogenic *E. coli.* Most others are believed to be important accessory genes for virulence of EAEC and other pathotypes [15,10,11,16,17].

Based on PCR and whole genome sequence analyses, it has previously been reported that SPATES are more common in EAEC and *Shigella* than other *E*. *coli* pathotypes [18,19]. There have also been reports associating SPATE genes with virulence in epidemiological studies [12, 20,21] and in an animal model [11]. We therefore sought to identify SPATE genes in EAEC that have been recovered from recent epidemiological studies in Nigeria and determine their association with disease. In the course of doing so, we found that rigorous methodology is required to correctly call SPATE genes in EAEC, and presumably other SPATE-rich genomes. Our observation led us to develop and validate suitable methodology and we found that earlier methods over-estimated the numbers and types of SPATEs in EAEC epidemiological strains sets. We were then able to evaluate the prevalence of SPATE-encoding genes among EAEC genomes derived from stool samples of children 0-5 years with or without diarrhoea and their association with disease.

## Methods

### Aim

This study was carried out to determine the prevalence of SPATE encoding-genes among EAEC genomes and their association with diarrhoea. *E. coli* strains were isolated from children 0-5 years with or without diarrhoea from two epidemiological studies conducted in Oyo State in Nigeria [22,23].

### Short read sequencing

Whole genome Illumina sequencing was carried out at the Wellcome Sanger Institute as part of the project titled “Pathogenic lineages of enteric bacteria in Nigeria”, as described by [22]. The reads have been deposited on ENA with the accession number PRJEB8667.

### Quality assurance and EAEC identification

Post-Sequencing quality control (QC) was first conducted on our paired end reads with FastQC (www.bioinformatics.babraham.ac.uk/projects/) and the results were aggregated with MultiQC [24]. Following the filtering using MultiQC report. Kraken [25] was used to assign genomic IDs to reads and Bracken [26] was used to quantify the genomic IDs assigned by Kraken. Reads that were reported as ‘failures’ with FastQC or have less than 90% reads assigned to E. coli species after Kraken genomic ID assignment and quantification of assigned genomic IDs were excluded from downstream analyses.

We defined EAEC as any genome having at least one of the following EAEC virulence genes–genes - *aggR*, *aar, aaiC, aap, aatA, aggA, aafA, agg3A,agg4A, agg5A, capU, air, afaD* and *eilA.* We heretofore screened 881 EAEC genomes derived from stool samples of children 0-5 years with or without diarrhoea for the presence of 23 SPATE-encoding genes (*sat*, *pet*, *vat*, *sigA*, *hbp*, *tsh*, *pic*, *eatA*, *eaaA*, *eaaC*, *sepA*, *espI*, *epeA*, *espP*, *espC*, *tleA*, *tagA*, tagB *sha*, *boa*, *picU*, *rpeA* and *crc1.* Furthermore, sequence reads were assembled using SPAdes genome assembler [27] and supplemented with Velvet genome assembler [28] was used instead. Following sequence assembly, contigs generated by the SPAdes [27] and Velvet [28] genome assemblers were checked for quality of genome assembly using CheckM [29] and supplemented with QUAST [30]. Genomes with more than 10% contamination rate from CheckM [29] results were excluded from downstream analyses.

### Phylogrouping and phylogenetic tree construction

To determine the phylogenetic groups of the *E. coli* in this study, reads that passed post-sequencing QC were assembled with SPAdes genome assembler [27]. Genome assemblies were then checked for quality using QUAST [30] and CheckM [29]. Ezclermont [31] was then used to determine the phylogenetic groups of the genomes. To construct phylogenetic trees, reads were mapped to the prototypical genome of EAEC 042 (FN65554) using a Wellcome Sanger Institute, Pathogen Informatics in-house tool – sh16_scripts (multiple_mappings_to_bam.py) [32]. *Escherichia fergusonnii* (NC_017626.1), was included in the reads as an outgroup. The mapping of reads to EAEC 042 (FN65554) was used to create multifasta alignment files of the reference genome and reads (including the outgroup genome). SNPs were called using snp-sites [33]. Phylogenetic trees were constructed using IQTree [34] using GTR+G model and trees were rooted with *Escherichia fergusonii*. The trees were viewed with Figtree [35], Phandango [36], iTOL [37] and Microreact [38].

### Annotation and feature identification

Assembled sequences were annotated with Prokka [39] and pangenome analysis was carried out using Roary [40] and Panaroo [41]. Multilocus Sequence Types (STs) of the assembled genomes were determined using ARIBA [42] and MLST (Achtman 7 loci) database [43]. Resistance genes were determined using ARIBA [42] and Resfinder database [44] and plasmid replicons were determined using ARIBA [42] and Plasmidfinder database [45]. In-silico serotyping was carried out using EcTyper [46] and in-silico pathotyping was carried out using ARIBA [42] and VirulenceFinder database [47].

### SPATE gene identification from short read sequence

In the first instance, SPATEs genes were detected using ARIBA [42] and Virulencefinder database [47]. For more stringent detection, we created a custom database [48] comprising SPATE-encoding genes of Virulencefinder database [47] Virulencefactor database [49] and some SPATEs whose function have been described, are not available on any of the two databases described above, but have been deposited on NCBI ([50]: *boa*, *sha*, *tagB*, *tagC*, *crc1*). For reference purposes, we included some SPATEs genes that are on either database and are also available on NCBI. Two SPATE-encoding genes (VFG033823-*sat* and VFG012922-*pic*) from virulence factor database were excluded from downstream analyses because the entries were about 10% of the estimated total gene length of the estimated lengths of *sat* and *pic*. Using this custom database, ARIBA was used for local assembly of SPATEs in our dataset. Interrupted, partial and fragmented assemblies were excluded from downstream analyses [51]. Where there is a discordance (partial vs complete/interrupted vs complete/fragmented vs complete) in the calls made by two or three databases, the call was adjudged to be correct based on any complete assembly of a specific gene from any one database and incorrect if a specific gene is not assembled completely. All data were analysed using simple statistics and chi square. A p value of <0.05 was considered statistically significant.

### SPATE identification from hybrid assemblies

Twenty-nine Illumina-sequenced genomes that carried SPATE-encoding genes (ARIBA/Virulencefinder database), were resequenced using Oxford Nanopore Technology (ONT), generating long reads thus— Genomic DNA libraries were prepared for long-read sequencing using the Oxford Nanopore Technologies (ONT) ligation sequencing chemistry with native barcoding and sequenced on a MinION Mk1C. Library preparation was performed on high-quality extracted genomic DNA using the Ligation Sequencing Kit (SQK-LSK109) in combination with the Native Barcoding Expansion Kit (EXP-NBD104), following the native barcoding protocol. Base-calling was performed using Guppy 5.0.17, with a minimum quality score of 8; reads that exceeded this threshold were retained for downstream analysis. These long reads were then matched with corresponding Illumina short reads and hybrid assembled using Unicycler [52]. Genomes were matched if any SPATE gene was detected in the genome, regardless of whether the detection was reported as “complete”, “fragmented” or “partial”. For Unicycler, all three modes (normal, conservative and strict) were used for assembly. Assemblies generated by the “normal” mode of Unicycler were selected for annotation because the “normal” mode has the lowest rate of mis-assembly [52]. Following assembly selection, assemblies were annotated with Bakta [53]. Contigs containing SPATE-encoding genes were viewed using Clinker [54], Proksee [55] and BRIG [56].

## Results

### Special precautions must be taken to avoid SPATE miscalls in SPATE-rich genomes

A total of 1,514 stool *E. coli* isolate genomes were included in this study of these, 881 met the definition of EAEC by virtue of possessing at least one of the following genes— *aggR*, *aar, aaiC, aap, aatA, aggA, aafA, agg3A,agg4A, agg5A, capU, air, afaD* and *eilA* [21]. The remaining 633 strains were non-EAEC. In the initial calls using only Virulencefinder database and ARIBA (Approach A), 1,434 SPATEs genes were identified with default settings, comprising *vat* (53, 4%), *pic* (109, 8%), *sepA* (328, 23%), *espI* (136,9%), *epeA* (47, 3%), *espC* (60, 4%), *espP* (54, 4%), *pet* (27, 2%), *sat* (463, 42%), *sigA* (80, 6%), *tsh* (29, 2%) and *eatA* (48, 3%). This initial analysis detected *sigA* in 54 EAEC from cases and 26 from controls, suggesting that the *sigA* gene is strongly associated with disease (the chi squared *p* value was 0.00001). SigA is a 103 kDa SPATE protein, earlier described on a chromosomal pathogenicity island in *Shigella* (Accession number AF600692, [57]). Attempts to determine the context of the encoding gene in two EAEC genomes from this study, LWD0045 and LLD0281 - both from children with diarrhoea - revealed that *sigA* was not present in the genome of either strain query. Instead, the Virulencefinder and BLAST matches to *sigA* arose from significant matches with chromosomal *pic* and *sat* genes in the highly conserved C-terminals (β-barrel regions), which are 67.89 % identical (Figure 1). Across the whole gene sequence, *sigA* is 39.7%% identical to *pic* and 54.73% to *sat* and, moreover, *pic* and *pet* show 40.06% identity with each other.

**Figure 1:**
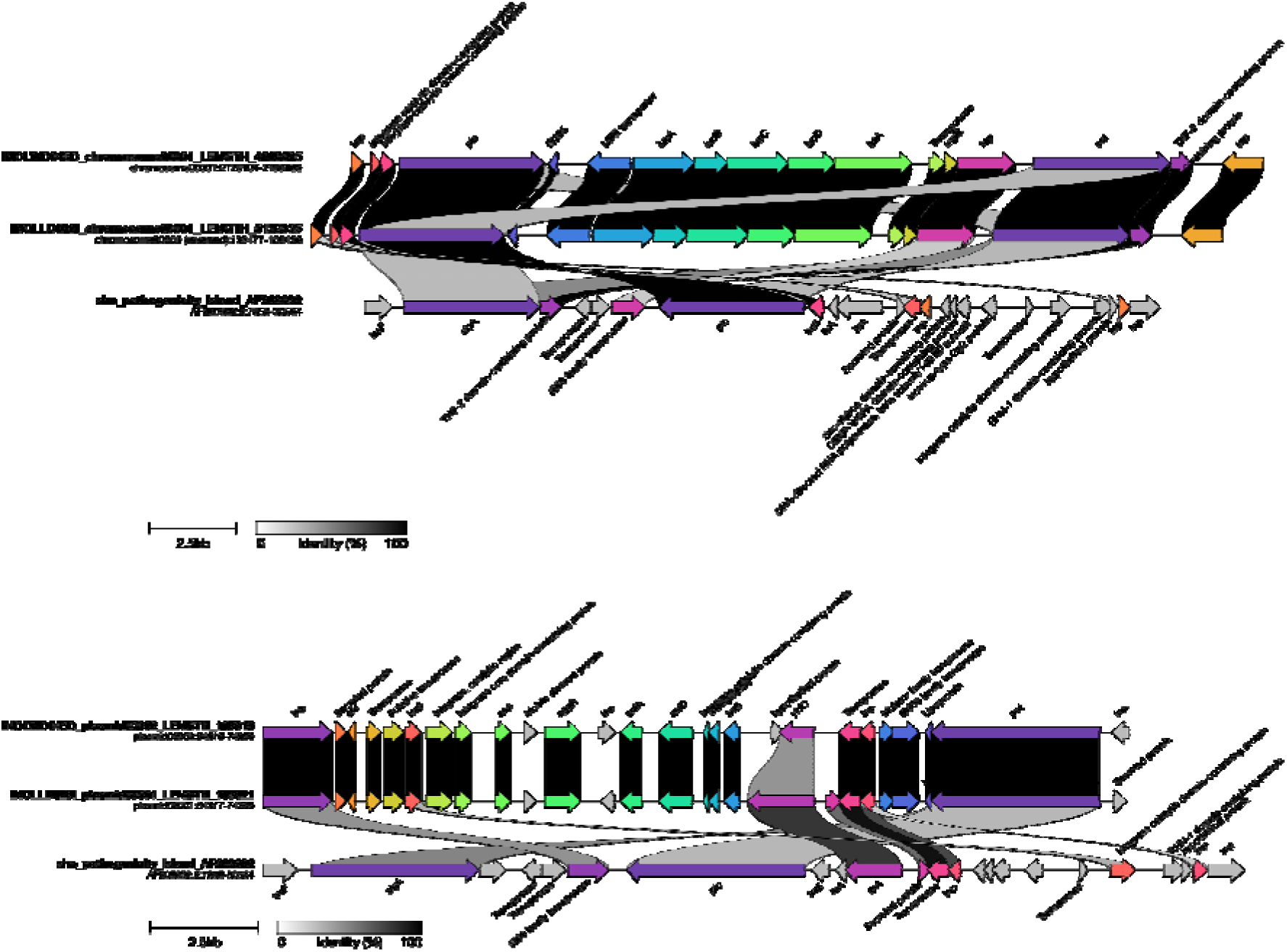
Similarity between different SPATES can lead to miscalls in virulence factor queries if coverage of the passenger region is not verified. Similarities between the *Shigella sigA* and (A) *pic* and *sat* on the chromosome and (B) *pet* on the plasmid of two EAEC strains from children with diarrhoea, LWD045D and LLD028I, that do not in fact carry the *sigA* gene.

Recognizing the risk of significant miscalls because the initial screen suggested that most EAEC carry multiple SPATEs, we developed a protocol to permit simple but accurate calls of SPATEs based on ARIBA [42]. First, we created a SPATE-only database which primarily consists of SPATE genes on Virulencefinder database, VirulenceFactor database and SPATE genes that are not on any of the two databases but have been characterised, annotated as SPATEs and are available on the NCBI. Incomplete SPATE genes, which have gene lengths that are a fraction (about 10%) of the estimated length of SPATE genes, were excluded from this new database. Second, using this custom database as reference, ARIBA was used for local assembly of SPATEs in our dataset. Interrupted, partial and fragmented assemblies were excluded from downstream analyses. When there is a discordance (partial vs complete/interrupted vs complete/fragmented vs complete) in the calls made by two or three databases, the call was adjudged to be correct based on any complete assembly of a specific gene from any one database. The initial Virulencefinder screen (Approach A—Table 1) identified 1,156 SPATE genes. However, when fragmented, interrupted or partial assemblies (hereafter referred to as incorrect calls) of SPATEs genes were removed, only 478 (41.3%) SPATEs genes were found by Approach B (Table 1). The other putative SPATE genes calls from Virulencefinder – 678 (58.7%) - appeared to be miscalls. All complete assemblies were referred to correct calls.

**Table 1:** Concordance/Disconcordance of SPATE-encoding genes detection for Illumina reads and hybrid assemblies.

| **ID** | **ARIBA/VFdb**<br>**Approach A** | **ARIBA/VFdb**<br>**(complete)** | **Bakta**<br>**(Unicycler_normal)**<br>**Approach B** | **Bakta**<br>**(Unicycler_conservative)** | **Bakta**<br>**(Unicycler_bold)** |
| --- | --- | --- | --- | --- | --- |
| LKD69F1 | *sepA*-partial; *espI*-partial; *sat*-partial | Nil | Nil | Nil | Nil |
| LKD69F2 | *sepA*-partial; *sat*-partial | Nil | Nil | Nil | Nil |
| LLD026A | *pic*-complete; *espI*-complete; *sepA*-partial; *epeA*-partial; *espP*-partial | *pic*; *espI*; *sat* | *pic*; *espI*; *sat* | *pic*; *espI*; *sat* | *pic*; *espI*; *sat* |
| LLD028I | *pic*-complete; *pet*-complete; *sat*-complete; *sigA*-partial | *pic*; *pet*; *sat* | *pic*; *pet*; *sat* | *pic*; *pet*; *sat* | *pic* ; *pet*; *sat* |
| LLD047A | *sepA*-partial; *sat*-partial | Nil | Nil | Nil | Nil |
| LLD106D | *pic*-complete; *pet*-complete; *sat*-complete | *pic*; *pet*; *sat* | *pic*; *pet*; *sat* | *pic*; *pet*; *sat* | *pic*; *pet*; *sat* |
| LLD52A2 | *sepA*-partial; *sat*-partial | Nil | Nil | Nil | Nil |
| LLH026C | *sepA*-partial; *sat*-partial | Nil | Nil | Nil | Nil |
| LLH342B | *sepA*-partial | Nil | Nil | Nil | Nil |
| LLH342D | *sepA*-partial | Nil | Nil | Nil | Nil |
| LWD034A | *pic*-partial; *sepA*-partial; *sat*-partial | Nil | Nil | Nil | Nil |
| LWD045E2 | *pic*-complete; *pet*-fragmented; *sat*-complete; *sigA*-partial | *pic*; *sat* | *pic*; *pet*; *sat* | *pic*; *pet*; *sat* | *pic*; *pet*; *sat* |
| MND044C | *sepA*-partial; *sat*-partial | Nil | Nil | Nil | Nil |
| MND60E | *sigA*-complete; *sepA*-partial; *espC*-partial, *sat*-partial | *sigA* | *sigA* | *sigA* | *sigA* |
| MND61B | *sepA*-partial; *sat*-partial; *sat*-partial | Nil | Nil | Nil | Nil |
| MND96E | *pic*-complete; *espI*-complete;<br>*sat*-complete; *sepA*-<br>fragmented; *epeA*-partial;<br>*espP*-partial | *pic; sat; espI* | *pic; sat; sepA; espI* | *pic; sat; sepA; espI* | *pic; sat; sepA; espI* |
| LWD45B | *pet*-complete; *espI*-partial;<br>*epeA*-partial; *espP*-partial;<br>*sigA*-partial | *pet* | *pet* | *pet* | *pet* |
| CHD063A | *sepA*-partial; *sat*-partial | Nil | Nil | Nil | Nil |
| CHD083J | *sepA*-partial; *sat*-partial | Nil | Nil | Nil | Nil |
| CHD277C | *sepA*-partial; *sat*-partial | Nil | Nil | Nil | Nil |
| LKD71A | *sepA*-partial; *sat*-partial | Nil | Nil | Nil | Nil |
| LLD025A | *sepA*-partial; *sat*-partial | Nil | Nil | Nil | Nil |
| LLD52A1 | *sepA*-partial; *sat*-partial | Nil | Nil | Nil | Nil |
| LLH190B | *sepA*-partial | Nil | Nil | Nil | Nil |
| LLH335B | *sepA*-partial; *sat*-partial | Nil | Nil | Nil | Nil |
| LLH342E | *sepA*-partial | Nil | Nil | Nil | Nil |
| LWD038A1 | *sepA*-partial; *sat*-partial | Nil | Nil | Nil | Nil |
| LWD038A2 | *sepA*-partial; *sat*-partial | Nil | Nil | Nil | Nil |
| MND81B | *sepA*-partial; *sat*-partial | Nil | Nil | Nil | Nil |

For ARIBA/VFdb, ARIBA was used for the detection of 12 SPATEs encoding genes found on the Virulencefinder database. ARIBA/VFdb (complete) contains a list of SPATE genes that have complete assemblies using ARIBA and Virulencefinder database. Unicycler\_normal, Unicycler\_conservative and Unicycler\_bold are the three assembly modes available in Unicycler hybrid assembler. Each of the assemblies from the three modes were annotated using Bakta and SPATEs found and confirmed by BLASTp were reported.

To validate this assessment, we long read sequenced 29 EAEC genomes on the Oxford Nanopore Technologies (ONT) platform and generated hybrid assemblies with the Illumina reads. As shown in Table 1, many of the ARIBA/Virulencefinder (Approach A) partial calls from short reads could not be located in hybrid assemblies. Comparing with SPATE calls from genomes that were hybrid assembled via Unicycler “Normal” mode (which has a lower rate of mis- assembly than “Bold” or “Conservative” [52]), we could observe that ARIBA/Virulencefinder short read calls that were complete were typically correct when we validated our results with reference SPATEs as shown in Table 1 and those that were partial were typically incorrect calls.

Correct calling varied with SPATE gene, with *pic*, *espI, sat, sigA* and *pet* always called correctly and *sepA* was often called incorrectly. Within this small dataset, we determined that the sensitivity and specificity of short read calls using ARIBA/Virulencefinder database and Bakta annotated “normal” Unicycler hybrid assemblies (Table 2). Accepting only complete calls “complete calls” from Virulencefinder output has high negative predictivity, although this protocol had low positive predictivity for *sepA* gene. The genomes of strains LKD69F, LLD52A and LWD038A were sequenced and hybrid-assembled twice (but only used once to compute test performance in Table 2) to verify that all methods supplied identical conclusions upon repetition.

**Table 2:** Sensitivity, Specificity, Accuracy, Positive Predictive Value and Negative Predictive Value for complete ARIBA/Virulencefinder SPATE calls vs calls generated from 29 hybrid assembly, annotated using Bakta and confirmed with BLASTp using reference SPATEs proteins.

| **SPATE** | **Sensitivity**<br>**(%)** | **Specificity**<br>**(%)** | **Accuracy**<br>**(%)** | **Positive Predictive Value**<br>**(%)** | **Negative Predictive Value**<br>**(%)** |
| --- | --- | --- | --- | --- | --- |
| *pic* | 100 | 100 | 100 | 100 | 100 |
| *espI* | 100 | 100 | 100 | 100 | 100 |
| *sat* | 100 | 100 | 100 | 100 | 100 |
| *pet* | 100 | 100 | 100 | 100 | 100 |
| *sigA* | 100 | 100 | 100 | 100 | 100 |
| *sepA* | 0 | 100 | 96.15 | - | 96.15 |

Having established that fragmented, partial and interrupted SPATEs assemblies could lead to miscalls of SPATE genes, we sought to first understand the reasons for these miscalls and second, understand the context of both miscalls and right calls in our dataset. We searched literature for protein sequences of SPATEs (Table 3) and compared with SPATEs in our dataset using BLASTp. Their percentage identity, coverage and length of alignment were compared.

**Table 3:** List and accession numbers of reference SPATEs proteins used for studying the context and cause of SPATEs miscalls.

| S/No | SPATEs | Reference | Source |
| --- | --- | --- | --- |
| 1. | SepA | CAA88252.1 | [58] |
| 2. | Pic | AAD23953.1 | [59] |
| 3. | Pet | AAC26634.1 | [60] |
| 4. | EspI | CAC39286.1 | [61] |
| 5. | Sat | AAG30168.1 | [62] |
| 6. | EspC | AAC44731.1 | [63] |
| 7. | Tsh | AAA24698.1 | [64] |
| 8. | EspP | CAA66144.1 | [65] |
| 9. | SigA | AAF67320.1 | [57] |

### The context for SPATES varies considerably among different EAEC strains

Querying short reads from our EAEC genomes using ARIBA and the Virulencefinder database yielded 25 presumptive *sepA* genes calls, one each from 25 SPATEs bearing genomes (Supplementary Table 1). However, running BLASTp using SepA proteins derived from hybrid assembled genomes and SepA reference protein (CAA88252.1), confirmed that only one of these genomes (MND96E) actually encoded SepA. Presumptive *sepA* genes calls for the other genomes were miscalls in which, as the BLASTp results showed, regions of 82% similarity of these SepA were only 133 amino acids of the conserved N-terminus (9% query coverage of the reference SepA protein (CAA88252.1)).

In phylogroup B1 strain MND96E, the *sepA* (98% identity with a query coverage of 90%) gene is borne on a 48,606bp IncFII plasmid which carries *bla*_TEM-1_ and *tetR*(A) (Figure 2). The plasmid carried what appeared to be a complete IncF conjugative system and no EAEC virulence genes except *sepA*. In this regard, the plasmid is a resistance plasmid and not an aggregative adherence virulence plasmid. The *sepA* gene was flanked by transposase and recombinase genes and had a G+C content of 48.79%, compared to 52.30% for the rest of the plasmid. Repetitive DNA around the *sepA* gene explains the ‘fragmented’ detection from short reads. The genome of strain MND96E carried three other plasmids, including an aggregative plasmid carrying *agg3ABCD*, *aggR*, *aar*, *aap*, *capU* EAEC-associated virulence genes.

**Fig 2:**
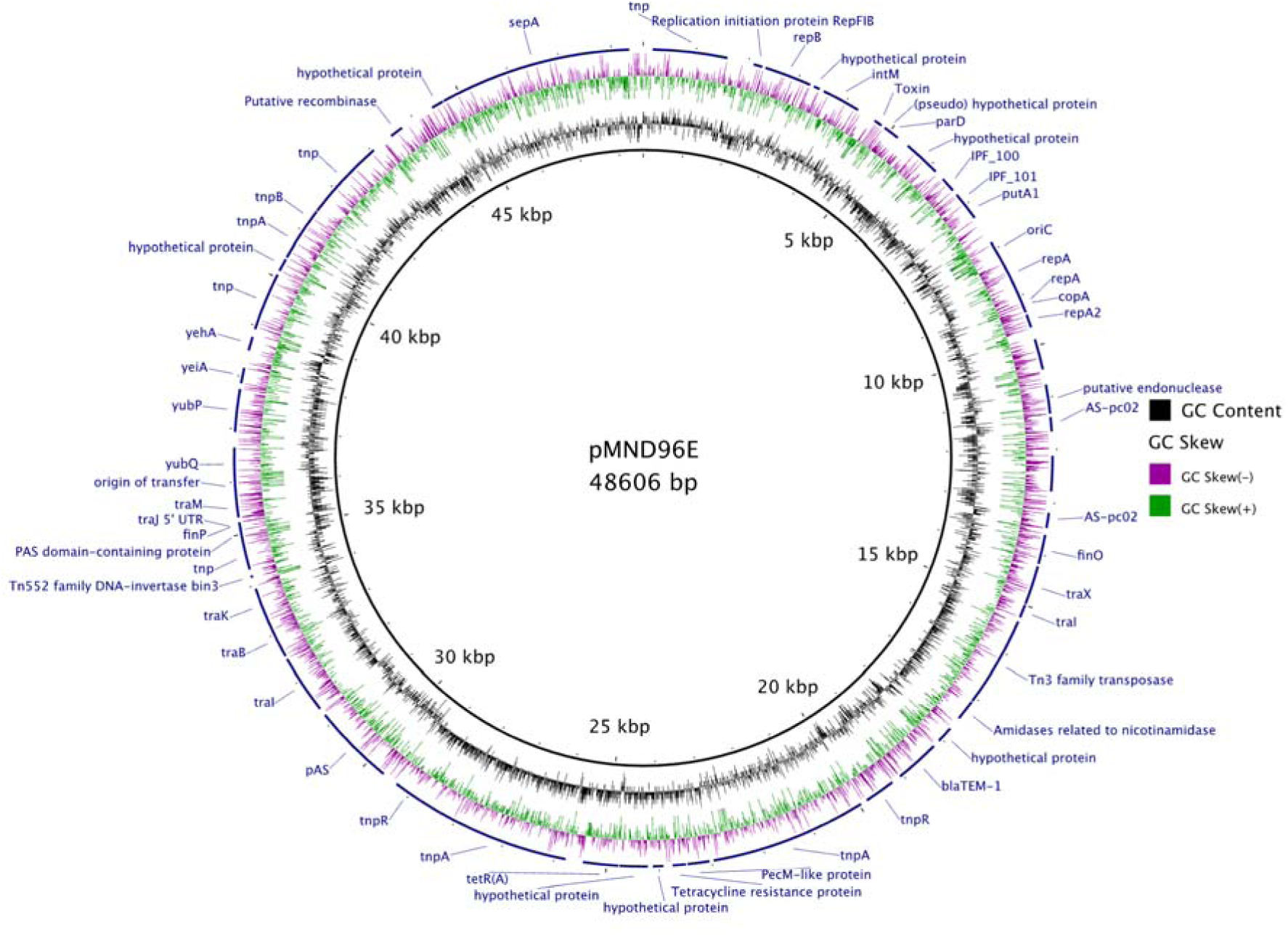
Map of the IncFIB resistance plasmid carrying *sepA* in the genome of EAEC strain MND96E

Figure 3 shows concentric maps of three *pet*-bearing plasmids recovered, one each, from LLD028I, LWD045E2 and LLD106D genomes aligned against the finished and closed pAA from reference EAEC strain 042 (Accession number FN554767.1). These three plasmids carry some EAEC-associated genes, including *aap*, *aar*, *aggR*, *aatA*, *capU* and AAF/I genes (*aafBCDA*).

**Fig. 3:**
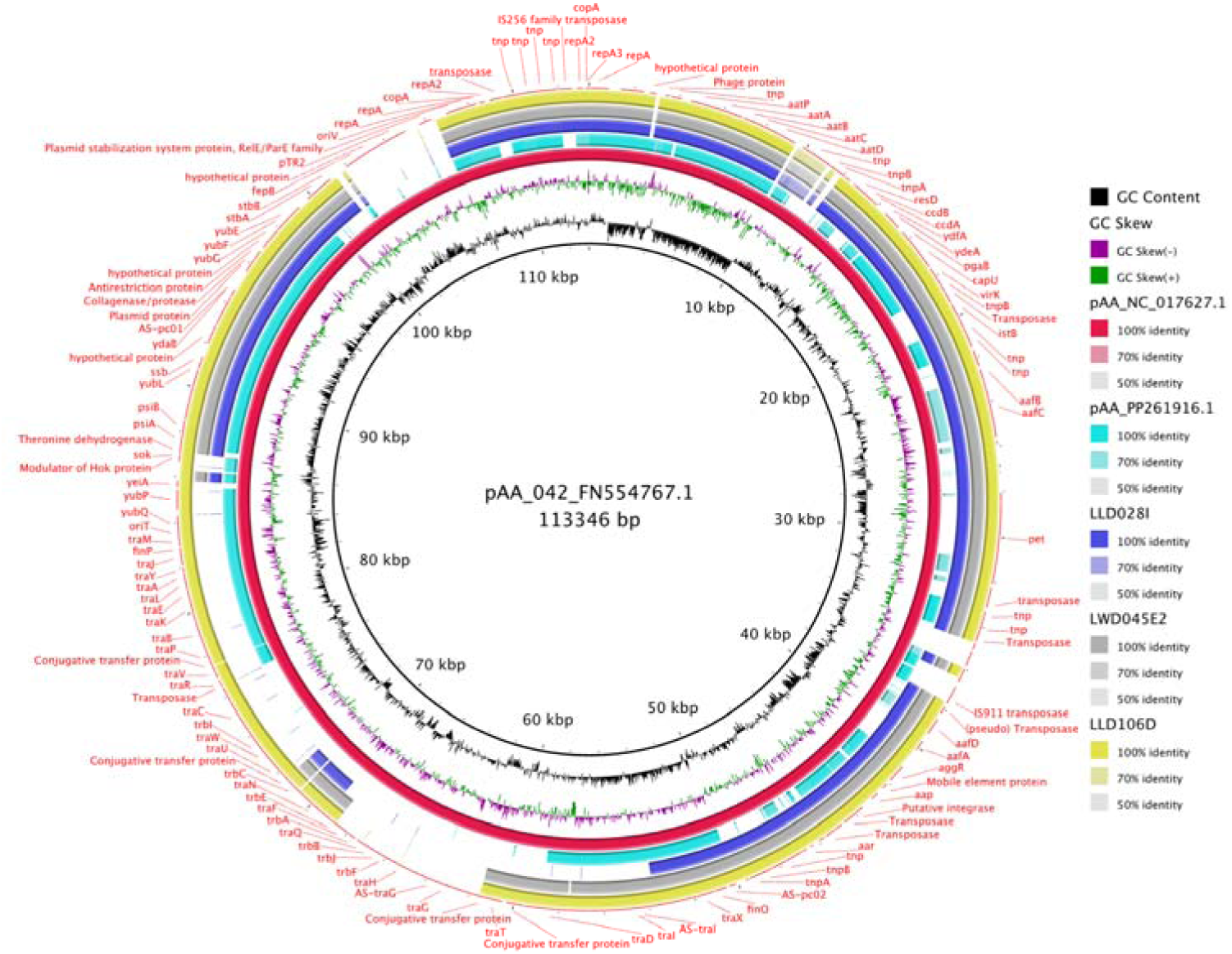
A circular map showing *pet*-bearing plasmids (pLLD028I-green ring, pLWD045E2-grey ring, pLLD106D-yellow ring) recovered from LLD028I, LWD045E2 and LLD106D genomes compared with pAA from FN554767.1. Other reference pAA plasmids, NC017627.1 (red) and PP261916.1 (green) were added to the ring.

Pet in LWD45B which also has 100% query coverage, but 99.85% percentage identity when compared with Pet reference using BLASTp (Supplementary Table 2). The others had 100% query coverage and percentage identity when compared with Pet reference.

Three presumptive *espC*-containing contigs (LWD034A, MND044C and MND96E) which also carried *sepA*, *aar*, *aggR* and *aap* are shown in Figure 4. As one includes *tra* genes and two of these three genomes carry AAF genes, which are typically plasmid-associated (*agg3ABCD* and *aggABCD*), the contigs are presumed originate from plasmids in EAEC strains MND96E and MND044C. However, comparison with reference EspC (AAC44731.1), uncovered that the *espC* ‘genes’ identified through ARIBA and the Virulencefinder database screening were actually limited coverage fragments of SPATEs and Immunoglobulin A1 protease (IgA1) Autotransporters N terminal conserved ends (Supplementary Table 2).

**Fig. 4:**
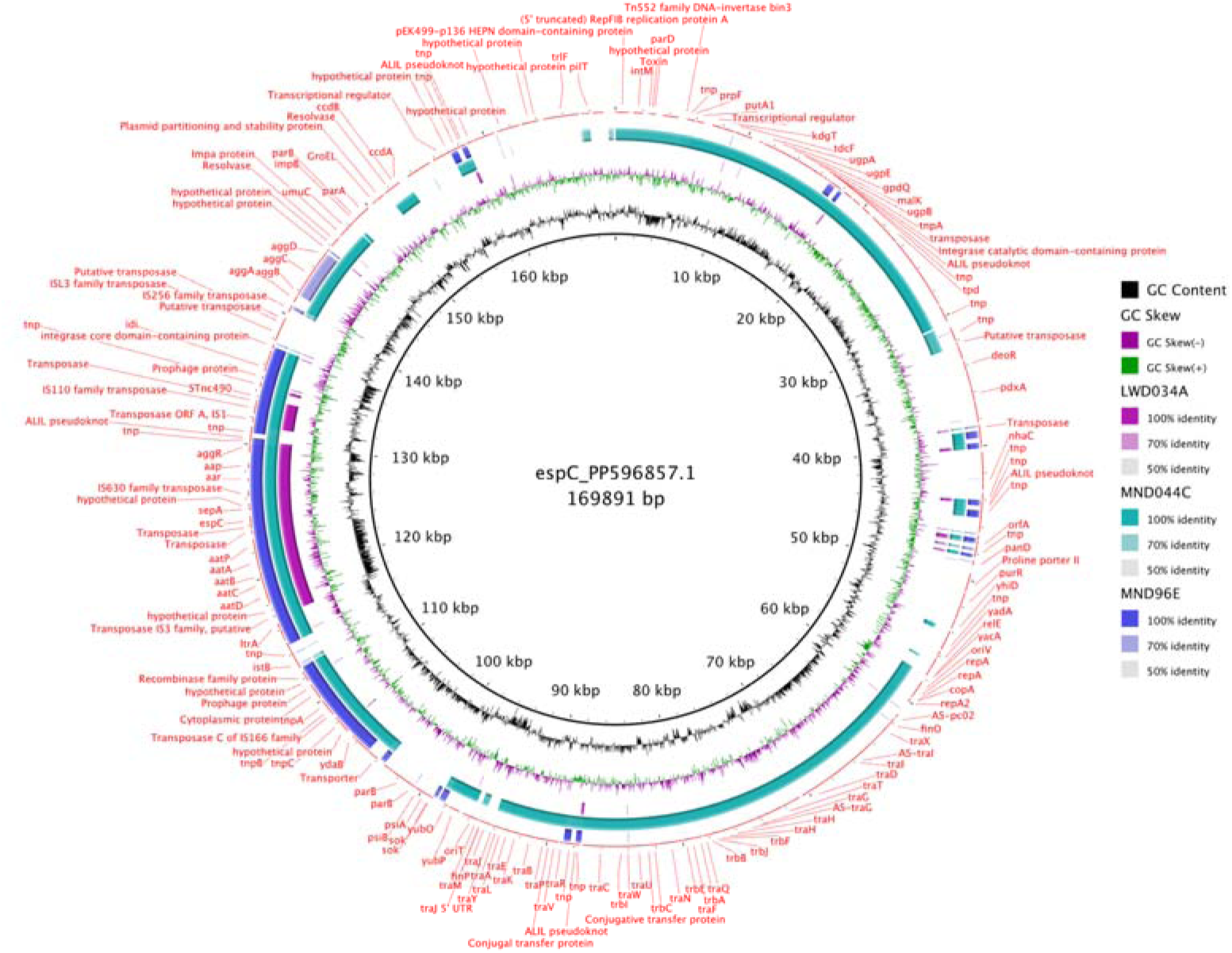
Comparison *espC* (PP596857.1) and 3 contigs bearing putative *espC* genes

Two contigs, one each from MND96E and LLD026A contained putative *espI* genes. When BLASTp was used to compare the EspI proteins from these two genomes using reference EspI protein (CAC39286.1), we found that the EspI had 100% query coverage and 99.05% percentage identity with the EspI reference (Supplementary Table 2). Context for *espI*, *tsh* and *espP* putative matches using BLASTp are detailed in the Supplementary Materials (Supplementary Table 2 and Supplementary Figure 1).

Three contigs, one each from LKD69F1, LKD69F2 and LWD034A carry putative *tsh* gene. Using BLAST similarity search against reference Tsh protein (AAA24698.1), we found that the putative *tsh* gene is identical to reference Tsh by only 40.7%. Further search using all SPATEs proteins in Table 2 revealed that of the nine reference SPATE proteins, the Tsh protein had the highest similarity (40.7%) with these three putative Tsh (Supplementary Table 3). Additional BLASTp search using reference proteins found on the GenBank gave a match (100% query coverage and 99.92% percentage identity) of an S6 family peptidase (WP_001531191.1). S6 family peptidases are similar to SPATEs and they belong to the same sub-group of the T5SS— classical autotransporters (Type Va) which consists of SPATEs, IgA1 proteases, subtilin-like serine proteases and Self-Associating Autotransporters (SAAT)

We found one contig in strain LWD034A carrying a putative espP gene. When we carried out a similarity search with EspP reference protein (CAA66144.1) using BLASTp, the EspP from strain LWD034A had 100% query coverage, but only 50.9% percentage identity with the reference protein. Supplementary Figure 2 shows a dot plot for the alignment of the reference EspP protein and the putative EspP found in LWD034A.

Following our observation that the putative EspP does not match reference EspP protein, we carried out BLASTp similarity search using all the reference SPATE proteins in Table 2. This gave a percentage identity of between 36.04% and 58.91%, which does not qualify for a match for any of the reference SPATE proteins. (Supplementary Table 4).

We observed that the *tsh* found in strains LKD69F1, LKD69F2 and LWD034A and the *espP* found in strain LWD034A all have autotransporter-like domains including IgA1 protease domain. We proceeded to carry out a similarity search between the putative Tsh and putative EspP proteins in these genomes. Our results show that while these putative SPATEs have 96% percentage coverage, their percentage identity was 35.07%. Interestingly, these putative SPATEs genes have amino acid lengths of 1309 aa (putative EspP) and 1279 aa (putative Tsh) which averages the number of amino acids that a typical SPATEs would have.

One *sigA* bearing contig from strain MND60E also carries *tnp* and *tra* genes required for genetic mobilisation and conjugation. Comparison of the putative SigA protein with reference SigA (AAF67320.1) showed that the SigA had 100% query coverage and 99.84%. Five presumptive *pic and sat* genes are carried on the same contig (contig 1) in genomes LLD106D, LWD045E2 and LLD028I and on different contigs in LLD026A and LLD028I. Comparison each of Pic and Sat from these genomes with Pic (AAD23953.1) and Sat (AAG30168.1) references respectively using BLASTp revealed that both the Pic and Sat proteins have 100% query coverage and at least 99% match with these references (Supplementary Table 2).

### While less abundant than earlier thought, EAEC strains from children in Nigeria frequently contain SPATE genes

Out of the 23 genes sought in the refined screen (Approach B), using ARIBA and our custom database, 19 SPATEs were detected in the same 881genome data set (compared to 12 using the original approach A. Four SPATEs (*rpeA*, *boa*, *hbp* and *picU*) were not found using either approach. Removing partial, interrupted and fragmented assemblies invalidated all earlier all calls for *espP*, *crc1* and *tleA* genes, so that 16 SPATEs were detected (*sat*, *pet*, *vat*, *sigA*, tsh, *pic*, *eatA*, *eaaA*, *eaaC*, *sepA*, *espI*, *epeA*, *espC*, *tagB*, *tagC* and *sha*).The calls for genes *espP*, *pic*, *sepA*, *espI*, *sat*, *sigA*, *tsh*, *espC*, and *tleA* differed significantly in the number and type of SPATEs detected using the two approaches (Figure 5).

**Fig. 5:**
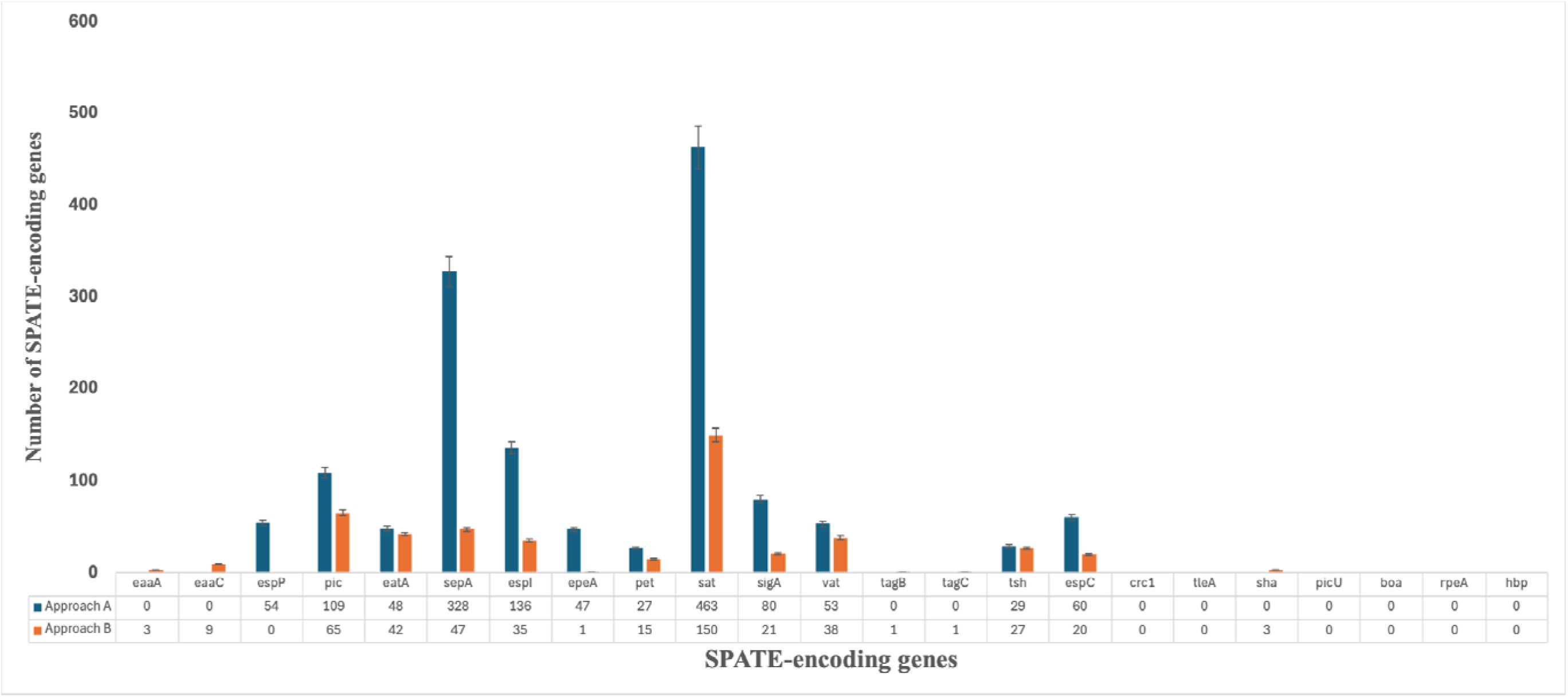
Variation in calls made for SPATEs between ARIBA/Virulencefinder database (Approach A) and refined ARIBA/SPATEs-custom database (Approach B). The absolute number of SPATE genes recovered using the two different approaches is also shown below the bars.

Using the refined (Approach B) protocol, 571 (64.8%) of 881 EAEC genomes screened had no SPATE gene, while 193 (21.9%) had only 1 SPATE gene compared to 248 (28.1%) and 168 (19.1%) respectively with the initial Approach A. Refined screening revealed that there were genomes carrying multiple SPATE genes – 68 (7.7%), 44 (5.0%) and 4 (0.5%) 2, 3 and 4 SPATE genes, respectively. However, these were fewer than those identified in the original, Approach A, with which we detected 271(30.7%), 113 (12.8%), 45 (5.1%), 20 (2.3%), 14 (1.6%) and 3 (0.3%) strains respectively carrying 2, 3, 4, 5, 6 and 7 SPATEs. Using the refined approach, 333 of the genomes carrying SPATEs were from children with diarrhoea and 548 from healthy children (multiple EAEC strains were isolated and sequenced from most children). The odds ratio for diarrhoea was 3.9 (*p*<0.0001) with one SPATE gene, 1.2 (*p*=0.6) with two SPATE genes, 1.4 (*p*=0.3) with three SPATE genes and 1.7 (*p*=0.6) with four SPATE genes (Figure 6). Thus, SPATE gene carriage was associated with disease, but multiple SPATEs were not associated with disease. While the presence of at least one SPATE was associated with disease, and therefore SPATEs might mark virulent EAEC, over half of the EAEC strains screened harboured no SPATE gene. Strains with no SPATEs (OR=0.7), showed no positive or negative association with diarrhoea.

**Fig. 6:**
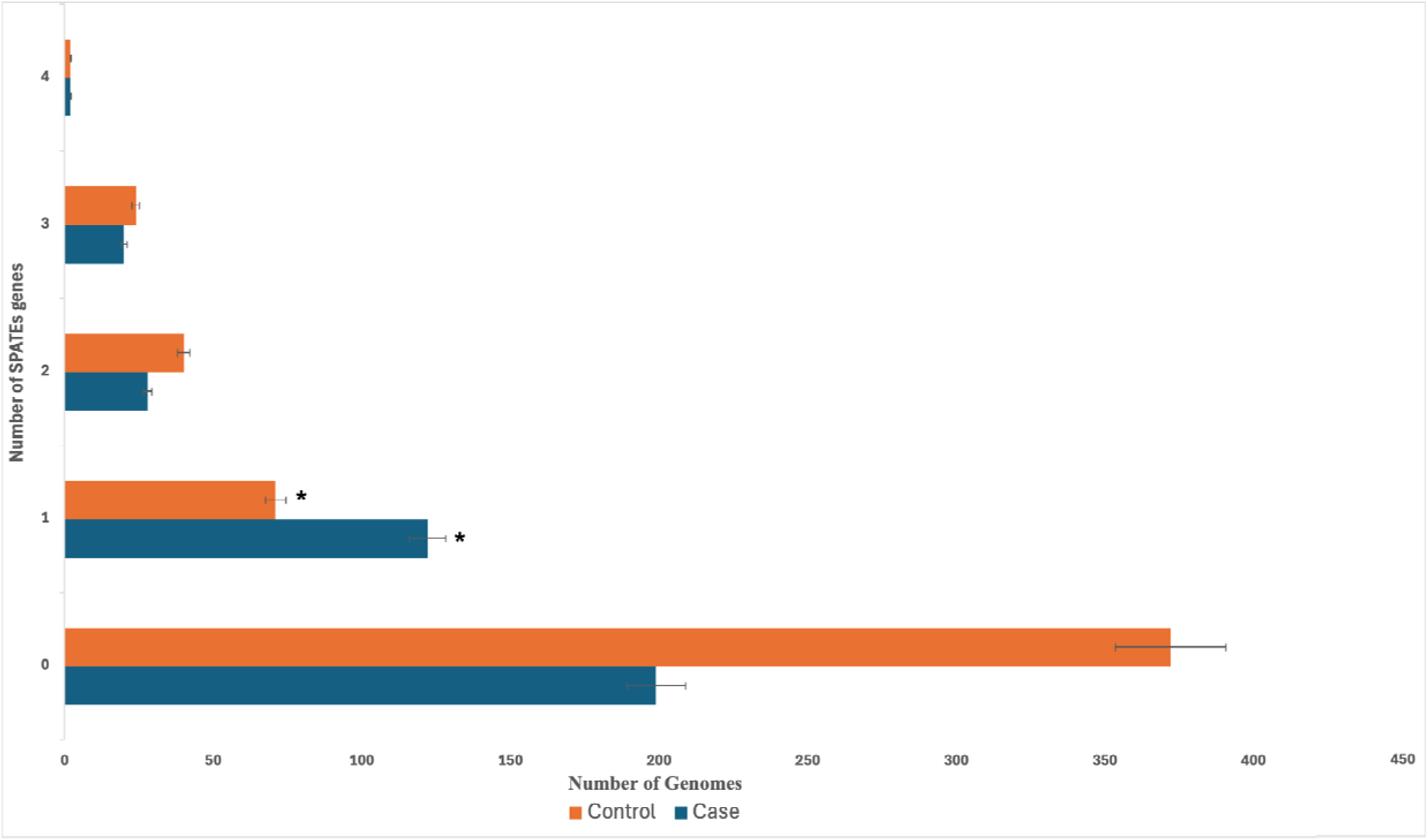
Occurrence of single or multiple SPATEs genes in EAEC genomes in isolates from children with diarrhoea (n= 333) and asymptomatic children (n=548). *(*p*<0.0001)

### SPATE gene miscalls in other datasets

As EAEC are reputed to be SPATE-rich [6,7,18,66,], typically carrying more than one SPATE in their genomes [18,21,67], we searched the literature for studies that had previously screened EAEC or other *E. coli* genomes for SPATE genes from Whole Genome Sequencing (WGS). We sought to determine whether others screening *E. coli* genomes for SPATES had encountered the same challenges we had and whether our Approach B would alter SPATE calls from their data sets. [68] carried out a study on *E. coli* isolated from bloodstream infections and reported that 50% (11/22) of their genomes carried SPATE genes with two carrying multiple SPATEs. When we re-interrogated their publicly available genomes (Accession PRJNA596854) using our Approach B methodology for SPATE gene detection, only one (4.5%) genome carried multiple SPATEs and their call for the *pic* gene was also incorrect (Table 3). As also shown in Table 4, [69] examined the genomic diversity of faecal non-diarrhoeagenic *E. coli* from children in sub-Saharan Africa and Asia. Of their 294 genomes, they reported that 119 (40.5%) carried at least one SPATE, while 175 (59.5%) bore no SPATE. When we used our modified, Approach B for SPATEs gene detection to rescreen the genomes retrieved from NCBI (Accession PRJNA611810), we discovered that their 68 calls for *espI* were all incorrect, the *pic* gene was always not called and there were mismatches between certain SPATE genes.

**Table 4:**
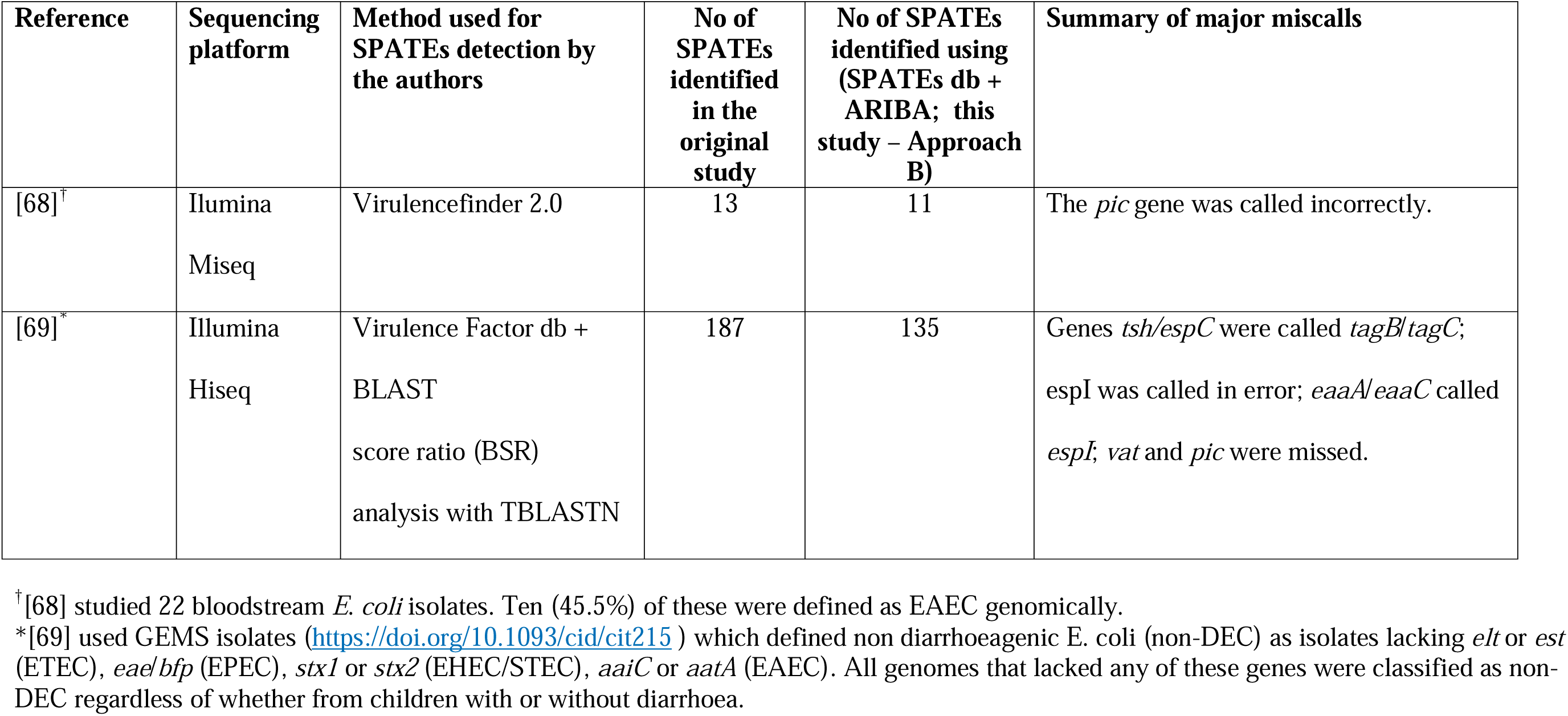
Summary of SPATE gene calls made by [68] and [69] and the results that we generated using their genomes and our refined methodology for SPATEs detection.

[21] and [67] papers were published without accession numbers for paired raw reads and therefore we could not retrieve the genomes used for their study and rescreen same using our modified approach.

### Rigorous screening demonstrates that SPATEs are abundant in the genomes of EAEC isolated in Nigeria

As shown in Figure 7, the most prevalent SPATE-encoding gene in our collection was *sat*, which was detected in 153(17.37%) EAEC, followed by *pic* 64 (7.26%). The *epeA*, *tagB* and *tagC* SPATE genes were least commonly detected, found in 1 (0.11%), 1 (0.11%) and 1 (0.11%) of the EAEC genomes, respectively. As shown in Figure 4, SPATE-positive EAEC were not distributed evenly across the EAEC phylogeny. Phylogroup A, B1 and C EAEC averaged about one SPATE per genome A(100/168), B1(165/158), C(9/11) while SPATE prevalence was lower in Phylogroups D(74/368) and F(14/52). EAEC bearing multiple SPATEs were most common in polygenetic group B2 (85 SPATEs /49 genomes), while SPATE genes were not present in genomes of EAEC belonging to phylogenetic groups E (0/20) and G (0/17). Undefined/cryptic phylogenetic group also carried many SPATEs genes (31/38).

**Fig. 7:**
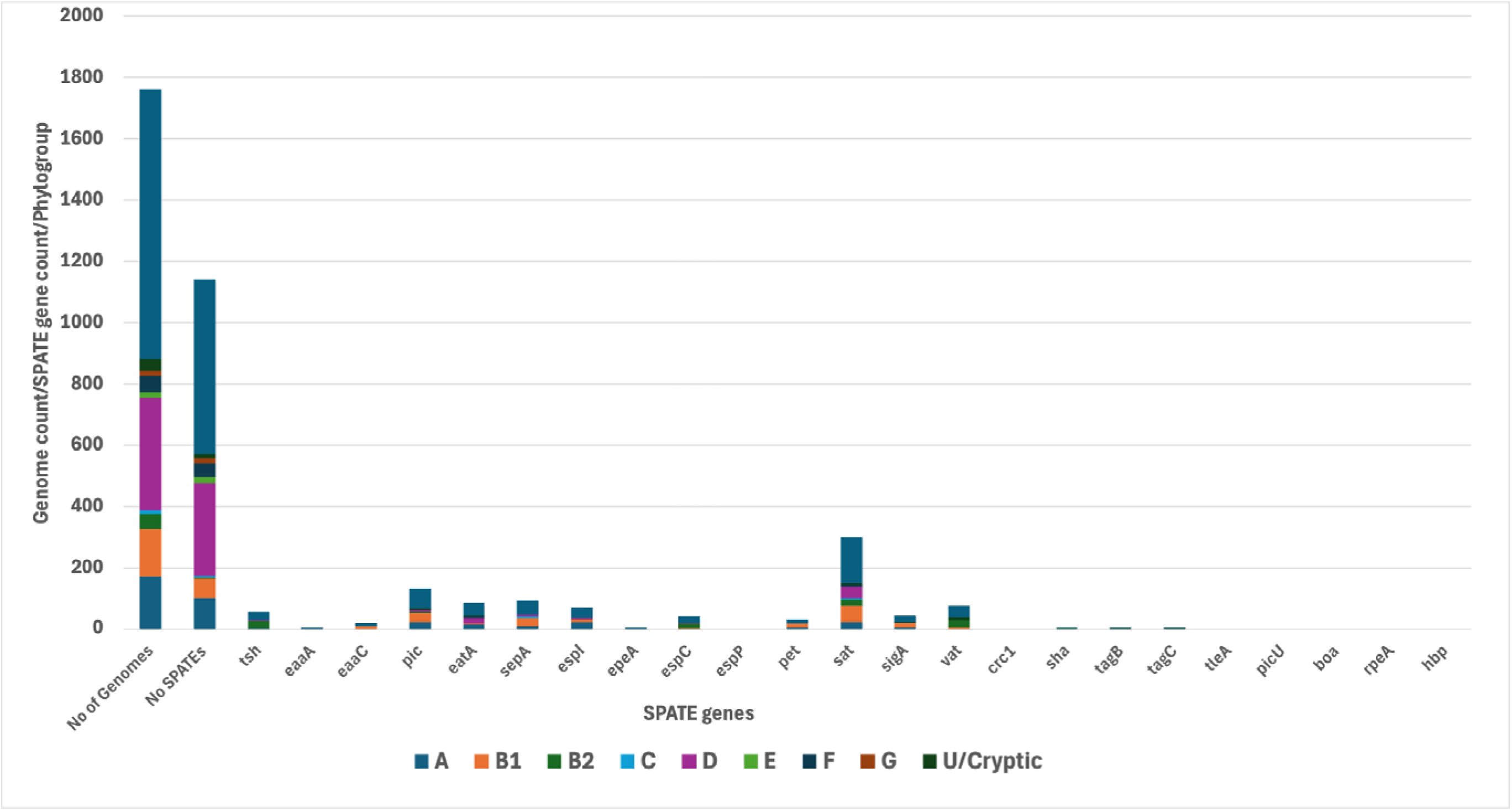
Abundance and distribution of SPATE genes by phylogenetic group

As shown in Table 5 shows the *pic* gene, detected in isolates from 40 (4.5%) children with diarrhoea and 25 (2.8%) healthy children, was significantly associated with diarrhoea (*p*=0.00004). The gene was also detected more frequently in isolates from children with diarrhoea (n=13, 1.5%) than healthy children (n=8, 0.9%) but the association was not significant after Bonferroni correction. However, the occurrence of *sigA* was associated with the carriage of multiple of SPATEs. Seventeen (81%) *sigA* positive strains also carried at least one other SPATE. Multiple SPATE gene carriage varies among SPATE-bearing strains (*pic*-85%; *eatA*-7%; *sepA*-38%; *espI*-74%; *pet*-73%; *sat*-52%; *vat*-50%; *espC*-90% and *tsh*-70%). SPATEs occurring with low frequencies *(epeA*, *eaaA*, *eaaC*, *sha*, *tagB* and *tagC*) all co-occur with at least one SPATE as follows: *epeA* (*espI*), *eaaA* (*sigA* and *sat*), *eaaC* (*sigA* and *sat*), *sha* (*pic*, *vat* and *espI*), *tagB* (*sat* and *tagC*) and *tagC* (*sat* and *tagB*).

**Table 5:**
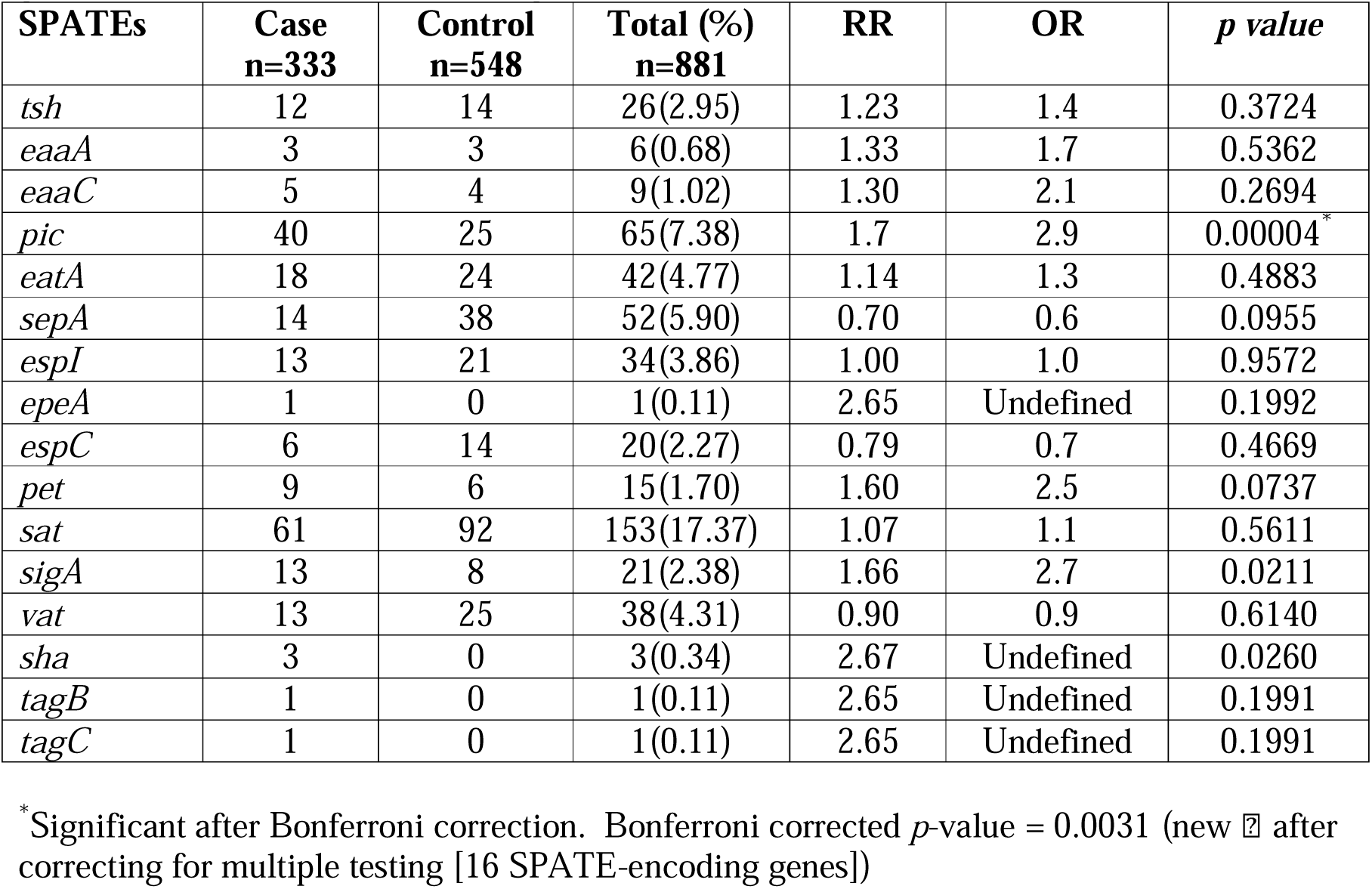
Association of SPATE-encoding genes with diarrhoea based on their presence in the genomes from isolates from four Nigerian case control studies.

Figure 8 shows a phylogenetic tree highlighting SPATEs genes detected in EAEC from cases and controls, as well as the number of SPATE genes detected per genome. The greatest diversity of SPATEs was seen in phylogroups B2, B1 and A, which contained most of the strains carrying multiple SPATEs. SPATEs that are commonly known to be chromosomal were infrequently detected among phylogroup D strains (the most common phylogroup), which typically carried 0 or 1 SPATE gene and for which *pet*, typically plasmid-borne, was the most common SPATE. There were few phylogroup E or G strains and none of them carried a SPATE gene. As shown in Figure 8 SPATE genes were evenly distributed between cases and controls in most phylogroups. However, they were associated with cases in less abundant phylogroups C (*p*=0.006) and F (*p*=0.01).

**Fig. 8:**
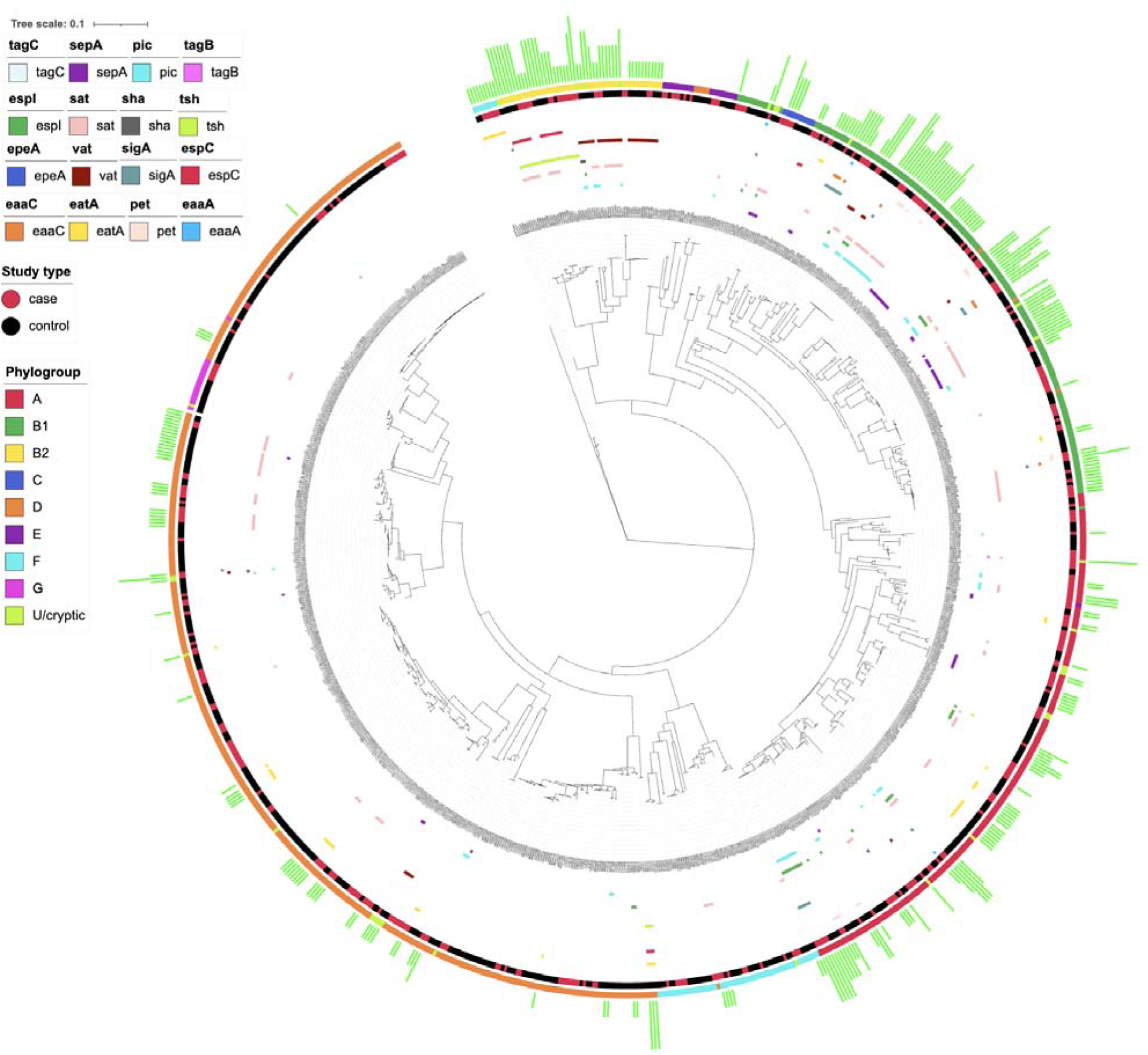
A maximum-likelihood core SNPs phylogenetic tree showing types and abundance SPATEs per EAEC genome and their phylogenetic groups

## Discussion

SPATES are important for *E. coli* pathogenesis and are among the best studied virulence factors in enteroaggregative *E. coli* (EAEC) [15,18,21,70,71,72]. Boisen et al observed that SPATEs are particularly common in EAEC genomes and reported that EAEC strains typically carry more than one SPATE gene [18]. SPATEs have additionally been proposed as critical virulence factors and potential markers to define the pathotype [21]. Our goal in this study was to determine the prevalence of SPATEs among EAEC and their association with diarrhoea using Virulencefinder database. We additionally developed and validated a refined protocol for the detection of SPATEs from WGS.

When we queried 881 short-read-sequenced EAEC genomes for SPATE genes with ARIBA/Virulencefinder database, we initially identified 1,156 partial, fragmented or complete presumptive SPATE-encoding genes. However, a more nuanced screen employing a custom database of SPATE-encoding genes on the Virulencefinder database [47], Virulencefactor database [49] and NCBI [50], found only 1,156 presumptive SPATEs. Removal of incomplete confirmed only 478 (41.3%) SPATEs were found. Results from our dataset demonstrate how indiscriminate use of a simple, often applied ARIBA/Virulencefinder database approach could lead to wrong and overestimated calls. We additionally applied our methodology to previous works that used Virulencefinder database/BLAST for the estimation of SPATE genes and got similar results, calling into question some of what is known about the molecular epidemiology of SPATE genes.

The earliest studies that highlighted over-representedness of SPATEs in EAEC genomes detected them by PCR. Andrade and others’ PCR-based study showed that of 193 EAEC in their study, 92(48%) carried multiple SPATEs and at least one SPATEs was detected in 137(71%) EAEC strains [66]. Boisen and her colleagues used a multiplex PCR to screen 55 EAEC strains for SPATE cytotoxins. They found 41(75%) of the strains carrying multiple SPATE genes, 11(20%) carrying single SPATE gene and 3(5%) without SPATEs [18]. While these earlier studies used PCR, PCR based methods on species as variable as *E. coli* typically have sensitivity or specificity shortfalls and WGS screens are also easier to execute for large gene sets, such as the 23 gene SPATE family. Therefore, more recent screens have justifiably leveraged WGS. In a study carried out by Petro et al to determine genotype associated with EAEC isolated from deployed military personnel with traveller’s diarrhoea using WGS data, five strains carried no SPATEs, 51(91%) carried multiple SPATEs [67]. Another study carried out by Boisen and others on isolates recovered from GEMS study using WGS data revealed that of the 97 EAEC strains that they screened for SPATE-encoding genes, 67(69%) carried at least one SPATE gene and 33(34%) carried multiple SPATE genes [21].

[68] conducted a study among blood stream infection hospitalised patients in Indonesia. The methods we devised in this study found that their *vat*, *sat* and *tsh* gene calls were correct but that *pic* was not. Similarly, applying our methodology to re-investigate Hazen et al’s genomes [69] found an over-estimation of SPATEs and multiple miscalls. Mismatches among SPATE gene sequences arise from conserved regions, which all or similar SPATEs share. While there is less homology among SPATE signal peptides and passenger domains, the linker regions are very highly conserved, and the C-terminal domain has a very high degree of amino acid homology. C-terminals of SPATEs are between 60% and 90% identical to one another [8]. EAEC strains, being SPATE-rich genomes, are particularly prone to this type of error as we have shown in this study. Additionally, some SPATEs are nearly identical, differing by only 2-4 residues [10]. These nearly identical SPATEs can be easily misidentified owing to their homology. For example, SPATE calls of *vat*, *tsh* and *hbp* in a single genome with percentage identity ranging from 70%-80%, ought to queried because *vat*, *tsh* and *hbp* all have at least 70% identity to each other. Double calling some SPATES also leads to an over-estimate of the number of SPATE genes each genome.

Homology among SPATEs accounts for most of the miscalls we uncovered, when a less refined approach is used. We additionally identified anthropomorphic contributions where databases contain indexed fragments of SPATE genes. For example, VFG033823-*sat* and VFG012922-*pic* indexed on the Virulencefactor database [49]. We additionally observed that there were incorrect annotations of SPATE genes by Bakta. These mis-annotations were responsible for the incorrect annotations of *espP*, *tsh* and other SPATE genes even in hybrid assemblies. The mis-annotation problems described above can be avoided if SPATEs calls use both high percentage identity (at least 99%) of nucleotides and high percentage coverage thresholds. In particular, coverage needs to be larger than the C-terminal beta barrel region of SPATEs, which is typically 750-900bp long.

## Conclusions

Correct identification of SPATE genes is important for understanding their molecular epidemiology within EAEC, their contribution to virulence in this little-understood pathotype as well as their potential as vaccine or therapeutic targets, if any. By developing and deploying a rigorous protocol for detecting and identifying SPATEs in *E*. *coli* genomes, we have laid the groundwork for understanding their role in pathogenicity. The results of our own study of EAEC from Nigeria support previous allegations that SPATEs are common in EAEC, but their prevalence is lower than earlier envisaged. We found SPATE association with diarrhoea overall, but importantly, SPATE distribution varied across the EAEC phylogeny, pointing to the likelihood that SPATEs may have different significances in different EAEC lineages. The *pet* gene, borne on a plasmid, is distributed largely without regard to phylogeny but most other highly prevalent SPATE genes, such as *pic, sat, sepA* were seen in some lineages but not others, *pic*, encoding a mucinase that has been shown to be important in EAEC pathogenicity [73,74] was found significantly more commonly in genomes of isolates from children with diarrhoea in our dataset. We however note that other SPATEs were associated with diarrhoea in some lineages but not others and multiple SPATEs were more commonly detected in phylogroups A, B1 and B2. Therefore, there may be important links to virulence within specific lineages of the EAEC pathotype. Future work needs to focus on understanding SPATE contribution to disease in different contexts, rather than presuming that these genes contribute identically in different bacterial and host backgrounds. This in turn will allow for researchers especially in regions where EAEC is endemic to identify truly pathogenic EAEC.

## List of Abbreviations

EAEC: enteroaggregative *Escherichia coli*
SPATEs: Serine Protease Autotransporters of Enterobacteriaceae
SAAT: Self-Associating Autotransporters
T5SS: Type V Secretion System

## Declarations

### Ethical approval and consent to participate

Ethical approval to conduct this research was granted by the University of Ibadan/University College Hospital Ibadan Research Ethical Committee (UI/UCH REC) with approval number UI/EC/15/0093.

### Consent for publication

Not Applicable

### Availability of data and materials

Primary data used for this study are available here: https://www.ebi.ac.uk/ena/browser/view/PRJEB8667. Third party data are available on NCBI (https://www.ncbi.nlm.nih.gov/) and their accession numbers are included in this published article. SPATEs database and protocols for detecting SPATEs are available on protocols.io and will cite this manuscript.

### Authors’ contributions

RAD conceptualised the idea, curated data, carried out formal analysis, investigation, visualization, original draft writing, review and editing. AOA conceptualised the idea, curated data, carried out formal analysis, investigation, visualization, review and editing. OAA curated data, carried out formal analysis, investigation, review and editing. BAT contributed to supervision, review and editing. BOO contributed to supervision, review and editing. NRT contributed to funding acquisition, supervision, review and editing. INO conceptualised the idea, acquired funding, curated data, supervised and performed original draft writing, review and editing.

All authors read and approved the final manuscript

## Acknowledgements

We thank Olabisi C Akinlabi, El-shama Q. Nwoko, Gabriel T Sunmonu, Stephen O. Bejide, Pelumi Daniel Adewole, Jola-Ade J. Ajiboye and David A Kwasi for technical and administrative assistance.

