## Supplemental Figures and Tables for "Refining the Serine Protease Autotransporters of Enterobacteriaceae (SPATE) gene detection in Enteroaggregative Escherichia coli genomes uncovers differential SPATE distribution by phylogeny"

**Supplementary Table 1:** Unvalidated Concordance/Disconcordance calls for SPATE-encoding genes detection for Illumina reads and hybrid assemblies

| **ID** | **ARIBA/VFdb** | **ARIBA/VFdb – complete only** | **Bakta (Unicycler_normal)** | **Bakta (Unicycler_conservative)** | **Bakta (Unicycler_bold)** |
| --- | --- | --- | --- | --- | --- |
| LKD69F1 | *sepA*-partial; *esp*I-partiai; *sat*-partial | Nil | *tsh* | *tsh* | *tsh* |
| LKD69F2 | *sepA*-partial; *sat*-partial | Nil | *tsh* | *tsh* | *tsh* |
| LLD026A | pic-complete; *espI*-complete; *sepA*-partial; *epeA*-partial; *espP*-partial | *pic; espI; sat* | *pic; espI; sat* | *pic; espI; sat* | *pic; espI; sat* |
| LLD028I | *pic*-complete; *pet*-complete; *sat*-complete; *sigA*-partial | *pic; pet; sat* | *pic; pet; sat* | *pic; pet; sat* | *pic ; pet; sat* |
| LLD047A | *sepA*-partial; *sat*-partial | Nil | *sepA* | *sepA* | *sepA* |
| LLD106D | *pic*-complete; *pet*-complete; *sat*-complete | *pic; pet; sat* | *pic; pet; sat* | *pic; pet; sat* | *pic; pet; sat* |
| LLD52A2 | *sepA*-partial; *sat*-partial | Nil | *sepA* | Nil | Nil |
| LLH026C | *sepA*-partial; *sat*-partial | Nil | *sepA* | Nil | *sepA* |
| LLH342B | *sepA*-partial | Nil | Nil | *sepA* | *sepA* |
| LLH342D | *sepA*-partial | Nil | *sepA* | *sepA* | *sepA* |
| LWD034A | *pic*-partial; *sepA*-partial; *sat*-partial | Nil | *espC; espP ; tsh* | *espC; espP; tsh* | *pic; sepA; sat; espC; espP; tsh* |
| LWD045E2 | *pic*-complete; *pet*-fragmented; *sat*-complete; *sigA*-partial | *pic; sat* | *pic; pet; sat* | *pic; pet; sat* | *pic; pet; sat* |
| MND044C | *sepA*-partial; *sat*-partial | Nil | *espC* | *espC* | *espC* |
| MND60E | *sigA*-complete; *sepA*-partial; *espC*-partial, *sat*-partial | *sigA* | *sigA* | *sigA* | *sigA* |
| MND61B | *sepA*-partial; *sat*-partial; *sat*-partial | Nil | Nil | *sepA* | Nil |
| MND96E | *pic*-complete; *espI*-complete; *sat*-complete; *sepA*-fragmented; *epeA*-partial; *espP*-partial | *pic; sat; espI* | *pic; sat; sepA; espI; espC* | *pic; sat; sepA; espI; espC* | *pic; sat; sepA; espI; espC* |
| LWD45B | *pet*-complete; *espI*-partial; *epeA*-partial; *espP*-partial; *sigA*-partial | *pet* | *espI; pet* | *espI; pet* | *espI; pet* |
| INOCHD063A | *sepA-*partial*; sat-*partial | Nil | Nil | Nil | Nil |
| INOCHD083J | *sepA-*partial*; sat-*partial | Nil | Nil | Nil | Nil |
| INOCHD277C | *sepA-*partial*; sat-*partial | Nil | Nil | Nil | Nil |
| INOLKD71A | *sepA-*partial*; sat-*partial | Nil | Nil | Nil | Nil |
| INOLLD025A | *sepA-*partial*; sat-*partial | Nil | Nil | Nil | Nil |
| INOLLD52A1 | *sepA-*partial*; sat-*partial | Nil | Nil | Nil | Nil |
| INOLLH190B | *sepA-*partial | Nil | Nil | Nil | Nil |
| INOLLH335B | *sepA-*partial*; sat-*partial | Nil | Nil | Nil | Nil |
| INOLLH342E | *sepA-*partial | Nil | Nil | Nil | Nil |
| INOLWD038A1 | *sepA-*partial*; sat-*partial | Nil | Nil | Nil | Nil |
| INOLWD038A2 | *sepA-*partial*; sat-*partial | Nil | Nil | Nil | Nil |
| INOMND81B | *sepA-*partial*; sat-*partial | Nil | Nil | Nil | Nil |

For ARIBA/VFdb, ARIBA was used for the detection of 12 SPATEs encoding genes found on the virulencefinder database. Unicycler_normal, unicycler_conservative and unicycler_bold are the three assembly mode available in unicycler hybrid assembler. Each of the assemblies from the three modes were annotated using Bakta and SPATEs found were reported in the genomes reported.

**Supplementary Table 2:** Summary of BLAST similarity search for putative SepA, Pet, EspC, Pic, Sat and EspI proteins

| **Genome** | **SPATE** | **Reference (Query)** | **Query Coverage(%)** | **% Identity** | **Alignment length** |
| --- | --- | --- | --- | --- | --- |
| MND96E | SepA | [CAA88252.1](https://www.ncbi.nlm.nih.gov/sites/entrez?cmd=Search&db=protein&term=CAA88252.1&dopt=GenBank) | 90 | 98.287 | 1226 |
| MND61B | SepA | [CAA88252.1](https://www.ncbi.nlm.nih.gov/sites/entrez?cmd=Search&db=protein&term=CAA88252.1&dopt=GenBank) | 9 | 81.746 | 126 |
| LLH342D | SepA | [CAA88252.1](https://www.ncbi.nlm.nih.gov/sites/entrez?cmd=Search&db=protein&term=CAA88252.1&dopt=GenBank) | 9 | 81.746 | 126 |
| LLH342B | SepA | [CAA88252.1](https://www.ncbi.nlm.nih.gov/sites/entrez?cmd=Search&db=protein&term=CAA88252.1&dopt=GenBank) | 9 | 81.746 | 126 |
| LLH026C | SepA | [CAA88252.1](https://www.ncbi.nlm.nih.gov/sites/entrez?cmd=Search&db=protein&term=CAA88252.1&dopt=GenBank) | 9 | 81.746 | 126 |
| LLD047A | SepA | [CAA88252.1](https://www.ncbi.nlm.nih.gov/sites/entrez?cmd=Search&db=protein&term=CAA88252.1&dopt=GenBank) | 9 | 81.746 | 126 |
| LLD52A2 | SepA | [CAA88252.1](https://www.ncbi.nlm.nih.gov/sites/entrez?cmd=Search&db=protein&term=CAA88252.1&dopt=GenBank) | 9 | 81.746 | 126 |
| LWD045E2 | Pet | [AAC26634.1](https://www.ncbi.nlm.nih.gov/sites/entrez?cmd=Search&db=protein&term=AAC26634.1&dopt=GenBank) | 100 | 100 | 1295 |
| LLD106D | Pet | [AAC26634.1](https://www.ncbi.nlm.nih.gov/sites/entrez?cmd=Search&db=protein&term=AAC26634.1&dopt=GenBank) | 100 | 100 | 1295 |
| LLD028I | Pet | [AAC26634.1](https://www.ncbi.nlm.nih.gov/sites/entrez?cmd=Search&db=protein&term=AAC26634.1&dopt=GenBank) | 100 | 100 | 1295 |
| LWD45B | Pet | [AAC26634.1](https://www.ncbi.nlm.nih.gov/sites/entrez?cmd=Search&db=protein&term=AAC26634.1&dopt=GenBank) | 100 | 99.85 | 1295 |
| MND044C | EspC | [AAC44731.1](https://www.ncbi.nlm.nih.gov/sites/entrez?cmd=Search&db=protein&term=AAC44731.1&dopt=GenBank) | 13 | 39.13 | 184 |
| LWD034A | EspC | [AAC44731.1](https://www.ncbi.nlm.nih.gov/sites/entrez?cmd=Search&db=protein&term=AAC44731.1&dopt=GenBank) | 13 | 39.13 | 184 |
| MND96E | EspC | [AAC44731.1](https://www.ncbi.nlm.nih.gov/sites/entrez?cmd=Search&db=protein&term=AAC44731.1&dopt=GenBank) | 17 | 38.59 | 184 |
| MND96E | Pic | [AAD23953.1](https://www.ncbi.nlm.nih.gov/sites/entrez?cmd=Search&db=protein&term=AAD23953.1&dopt=GenBank) | 100 | 99.78 | 1372 |
| LLD026A | Pic | [AAD23953.1](https://www.ncbi.nlm.nih.gov/sites/entrez?cmd=Search&db=protein&term=AAD23953.1&dopt=GenBank) | 100 | 99.78 | 1372 |
| LLD028I | Pic | [AAD23953.1](https://www.ncbi.nlm.nih.gov/sites/entrez?cmd=Search&db=protein&term=AAD23953.1&dopt=GenBank) | 100 | 99.71 | 1372 |
| LWD045E2 | Pic | [AAD23953.1](https://www.ncbi.nlm.nih.gov/sites/entrez?cmd=Search&db=protein&term=AAD23953.1&dopt=GenBank) | 100 | 99.71 | 1372 |
| LLD106D | Pic | [AAD23953.1](https://www.ncbi.nlm.nih.gov/sites/entrez?cmd=Search&db=protein&term=AAD23953.1&dopt=GenBank) | 100 | 99.71 | 1372 |
| MND96E | Sat | [AAG30168.1](https://www.ncbi.nlm.nih.gov/sites/entrez?cmd=Search&db=protein&term=AAG30168.1&dopt=GenBank) | 100 | 99.38 | 1295 |
| LLD026A | Sat | [AAG30168.1](https://www.ncbi.nlm.nih.gov/sites/entrez?cmd=Search&db=protein&term=AAG30168.1&dopt=GenBank) | 100 | 99.31 | 1295 |
| LLD028I | Sat | [AAG30168.1](https://www.ncbi.nlm.nih.gov/sites/entrez?cmd=Search&db=protein&term=AAG30168.1&dopt=GenBank) | 100 | 99.38 | 1295 |
| LWD045E2 | Sat | [AAG30168.1](https://www.ncbi.nlm.nih.gov/sites/entrez?cmd=Search&db=protein&term=AAG30168.1&dopt=GenBank) | 100 | 99.38 | 1295 |
| LLD106D | Sat | [AAG30168.1](https://www.ncbi.nlm.nih.gov/sites/entrez?cmd=Search&db=protein&term=AAG30168.1&dopt=GenBank) | 100 | 99.38 | 1295 |
| LLD026A | EspI | [CAC39286.1](https://www.ncbi.nlm.nih.gov/sites/entrez?cmd=Search&db=protein&term=CAC39286.1&dopt=GenBank) | 100 | 99.05 | 1363 |
| MND96E | EspI | [CAC39286.1](https://www.ncbi.nlm.nih.gov/sites/entrez?cmd=Search&db=protein&term=CAC39286.1&dopt=GenBank) | 100 | 99.05 | 1363 |

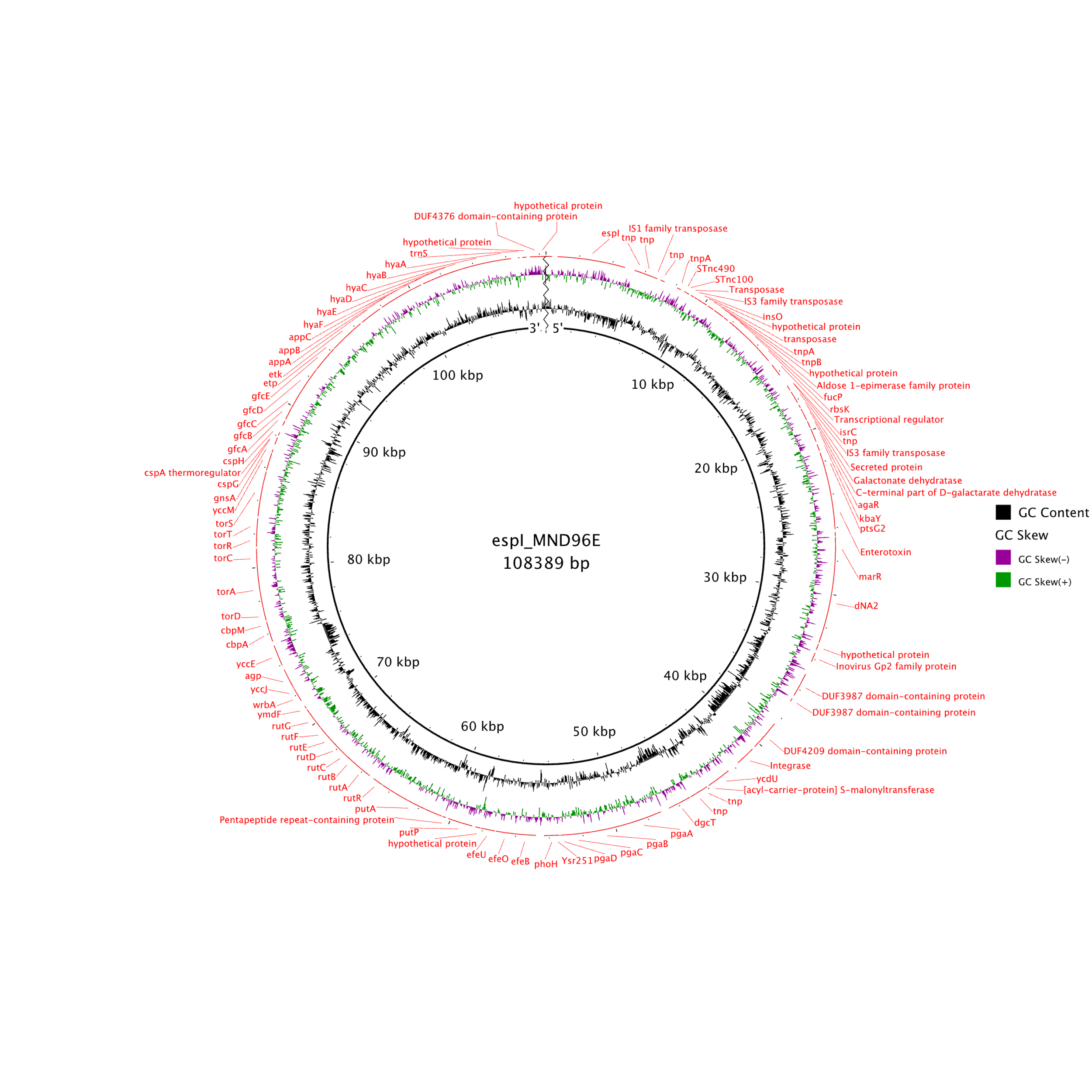
A. B

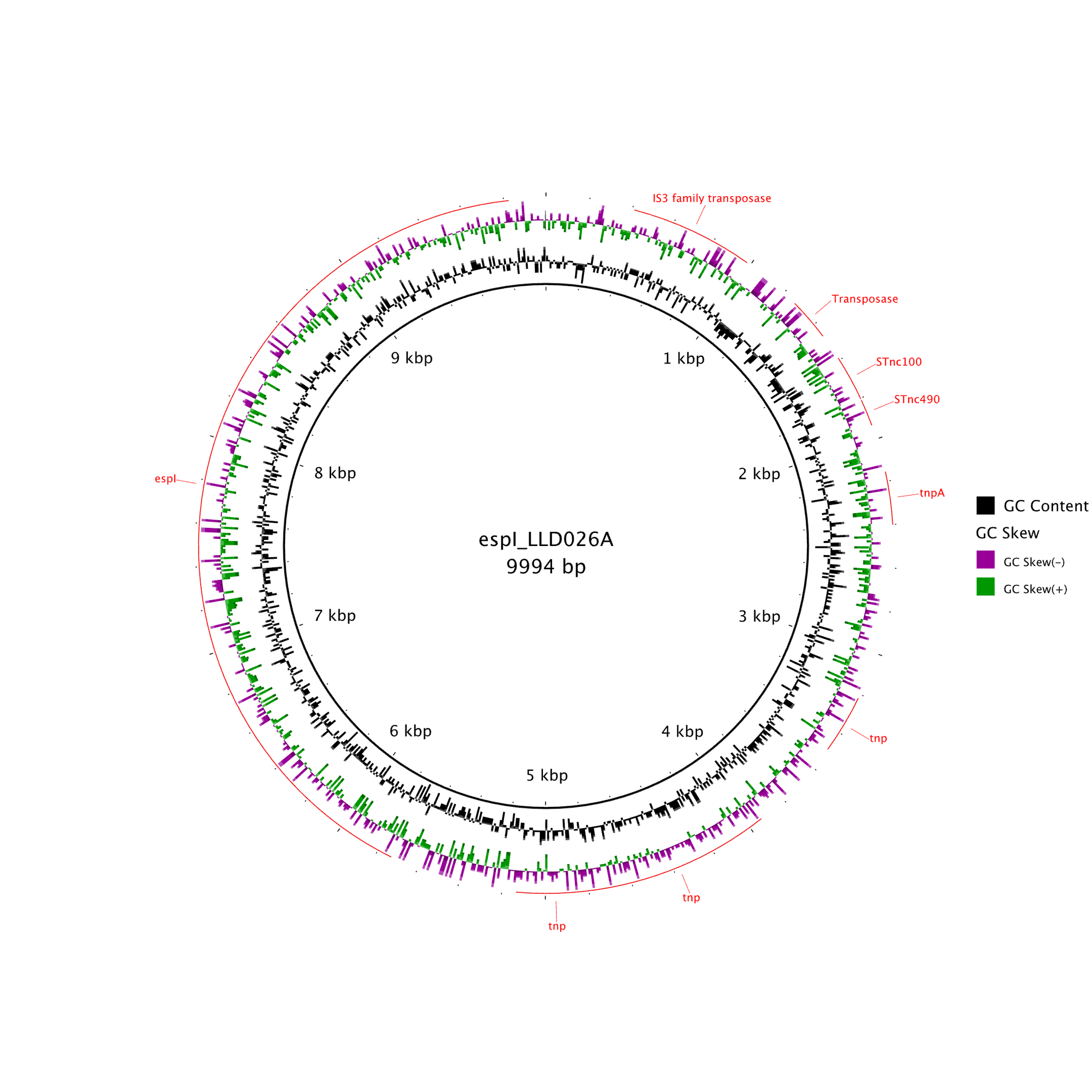

Supplementary Figure 1: Context of *espI* genes in contigs bearing the gene in (A.) strain MND96E and (B.) strain LLD026A

**Supplementary Table 3:** BLASTp results for putative Tsh from LKD69F1, LKD69F2, LWD034A and Reference SPATE proteins

| **SPATE** | **Accession Number** | **Query Length** | **Query Coverage(%)** | **% Identity** | **Alignment length** |
| --- | --- | --- | --- | --- | --- |
| SepA  LKD69F1  LKD69F2  LWD034A | [CAA88252.1](https://www.ncbi.nlm.nih.gov/sites/entrez?cmd=Search&db=protein&term=CAA88252.1&dopt=GenBank) | 1366  1366  1366 | 96  96  96 | 38.39  38.39  38.39 | 1279  1279  1279 |
| Pic  LKD69F1  LKD69F2  LWD034A | [AAD23953.1](https://www.ncbi.nlm.nih.gov/sites/entrez?cmd=Search&db=protein&term=AAD23953.1&dopt=GenBank) | 1372  1372  1372 | 92  92  92 | 40.36  40.36  40.36 | 1279  1279  1279 |
| Pet  LKD69F1  LKD69F2  LWD034A | [AAC26634.1](https://www.ncbi.nlm.nih.gov/sites/entrez?cmd=Search&db=protein&term=AAC26634.1&dopt=GenBank) | 1295  1295  1295 | 97  97  97 | 33.56  33.56  33.56 | 1279  1279  1279 |
| EspI  LKD69F1  LKD69F2  LWD034A | [CAC39286.1](https://www.ncbi.nlm.nih.gov/sites/entrez?cmd=Search&db=protein&term=CAC39286.1&dopt=GenBank) | 1363  1363  1363 | 98  98  98 | 38.10  38.10  38.10 | 1279  1279  1279 |
| Sat  LKD69F1  LKD69F2  LWD034A | [AAG30168.1](https://www.ncbi.nlm.nih.gov/sites/entrez?cmd=Search&db=protein&term=AAG30168.1&dopt=GenBank) | 1295  1295  1295 | 97  97  97 | 32.86  32.86  32.86 | 1279  1279  1279 |
| EspC  LKD69F1  LKD69F2  LWD034A | [AAC44731.1](https://www.ncbi.nlm.nih.gov/sites/entrez?cmd=Search&db=protein&term=AAC44731.1&dopt=GenBank) | 1306  1306  1306 | 95  95  95 | 34.74  34.74  34.74 | 1279  1279  1279 |
| Tsh  LKD69F1  LKD69F2  LWD034A | [AAA24698.1](https://www.ncbi.nlm.nih.gov/sites/entrez?cmd=Search&db=protein&term=AAA24698.1&dopt=GenBank) | 1377  1377  1377 | 98  98  98 | 40.7  40.7  40.7 | 1279  1279  1279 |
| EspP  LKD69F1  LKD69F2  LWD034A | [CAA66144.1](https://www.ncbi.nlm.nih.gov/sites/entrez?cmd=Search&db=protein&term=CAA66144.1&dopt=GenBank) | 1300  1300  1300 | 97  97  97 | 33.36  33.36  33.36 | 1279  1279  1279 |
| sigA  LKD69F1  LKD69F2  LWD034A | AAF67320.1 | 1285  1285  1285 | 95  95  95 | 33.15  33.15  33.15 | 1279  1279  1279 |

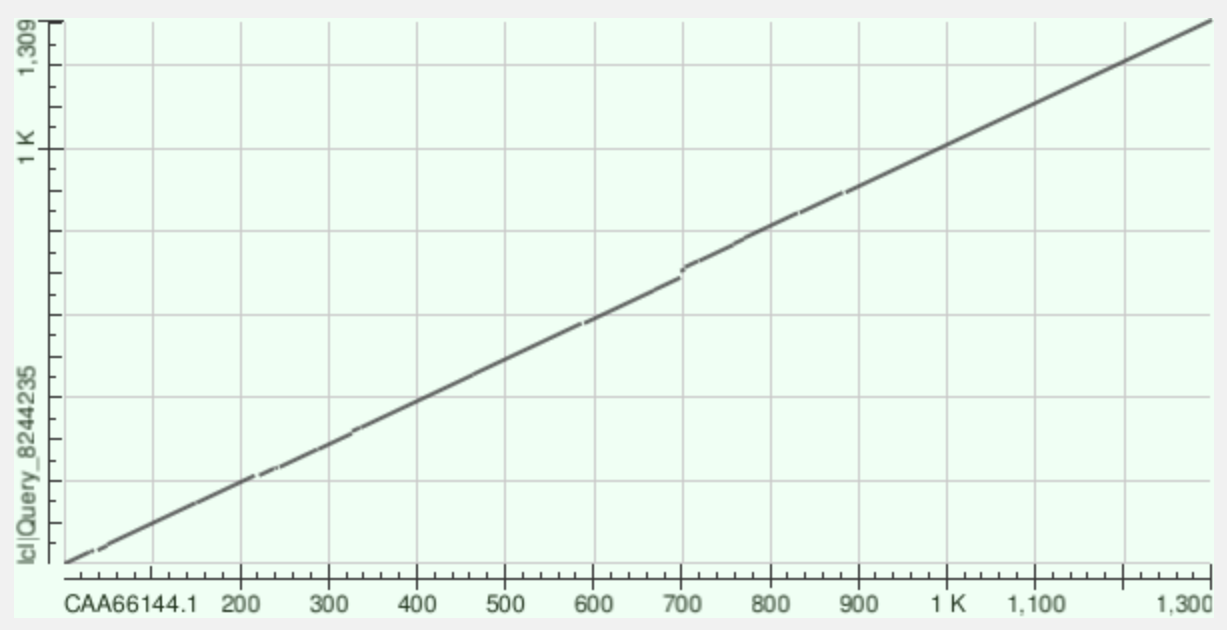

**Supplemntary Figure 2:** A dot plot showing the alignment of putative EspP from LWD034A on the y-axis and reference EspP (CAA66144.1) on the x-axis. EspP from LWD034A is 1309 amino acid long and reference EspP(CAA66144.1) is 1300 amino acid long

**Supplementary Table 4:** BLASTp results for putative EspP from LWD034A and Reference SPATE proteins

| **SPATE** | **Accession Number** | **Query Length** | **Query Coverage(%)** | **% Identity** | **Alignment length** |
| --- | --- | --- | --- | --- | --- |
| SepA | [CAA88252.1](https://www.ncbi.nlm.nih.gov/sites/entrez?cmd=Search&db=protein&term=CAA88252.1&dopt=GenBank) | 1366 | 94 | 49.25 | 1309 |
| Pic | [AAD23953.1](https://www.ncbi.nlm.nih.gov/sites/entrez?cmd=Search&db=protein&term=AAD23953.1&dopt=GenBank) | 1372 | 100 | 51.45 | 1309 |
| Pet | [AAC26634.1](https://www.ncbi.nlm.nih.gov/sites/entrez?cmd=Search&db=protein&term=AAC26634.1&dopt=GenBank) | 1295 | 100 | 50.53 | 1309 |
| EspI | [CAC39286.1](https://www.ncbi.nlm.nih.gov/sites/entrez?cmd=Search&db=protein&term=CAC39286.1&dopt=GenBank) | 1363 | 100 | 43.60 | 1309 |
| Sat | [AAG30168.1](https://www.ncbi.nlm.nih.gov/sites/entrez?cmd=Search&db=protein&term=AAG30168.1&dopt=GenBank) | 1295 | 100 | 51.40 | 1309 |
| EspC | [AAC44731.1](https://www.ncbi.nlm.nih.gov/sites/entrez?cmd=Search&db=protein&term=AAC44731.1&dopt=GenBank) | 1306 | 100 | 58.91 | 1309 |
| Tsh | [AAA24698.1](https://www.ncbi.nlm.nih.gov/sites/entrez?cmd=Search&db=protein&term=AAA24698.1&dopt=GenBank) | 1377 | 100 | 36.04 | 1309 |
| EspP | [CAA66144.1](https://www.ncbi.nlm.nih.gov/sites/entrez?cmd=Search&db=protein&term=CAA66144.1&dopt=GenBank) | 1300 | 100 | 50.90 | 1309 |
| SigA | AAF67320.1 | 1285 | 100 | 52.79 | 1309 |
